# A Primer on Modeling and Measurement of Signaling Outcomes Affecting Decision Making in Cells: Methods for Determining Optimal and Incorrect Outcomes in Noisy Biochemical Dynamics

**DOI:** 10.1101/2019.12.21.885491

**Authors:** Mustafa Ozen, Tomasz Lipniacki, Andre Levchenko, Effat S. Emamian, Ali Abdi

**Author notes:** Corresponding Author: Ali Abdi.

## Abstract

Characterization of decision makings in a cell in response to received signals is of high importance for understanding how cell fate is determined. The problem becomes multi-faceted and complex when we consider cellular heterogeneity and dynamics of biochemical processes. In this paper, we present a unified set of decision-theoretic and statistical signal processing methods and metrics to model the precision of signaling decisions, given uncertainty, using single cell data. First, we introduce erroneous decisions that may result from signaling processes, and identify false alarm and miss event that are associated with such decisions. Then, we present an optimal decision strategy which minimizes the total decision error probability. The optimal decision threshold or boundary is determined using the maximum likelihood principle that chooses the hypothesis under which the data are most probable. Additionally, we demonstrate how graphing receiver operating characteristic curve conveniently reveals the trade-off between false alarm and miss probabilities associated with different cell responses. Furthermore, we extend the introduced signaling outcome modeling framework to incorporate the dynamics of biochemical processes and reactions in a cell, using multi-time point measurements and multi-dimensional outcome analysis and decision making algorithms. The introduced multivariate signaling outcome modeling framework can be used to analyze several molecular species measured at the same or different time instants. We also show how the developed binary outcome analysis and decision making approach can be extended to include more than two possible outcomes. To show how the overall set of introduced models and methods can be used in practice and as an example, we apply them to single cell data of an intracellular regulatory molecule called Phosphatase and Tensin homolog (PTEN) in a p53 system, in wild-type and abnormal, e.g., mutant cells. These molecules are involved in tumor suppression, cell cycle regulation and apoptosis. The unified signaling outcome modeling framework presented here can be applied to various organisms ranging from simple ones such as viruses, bacteria, yeast, and lower metazoans, to more complex organisms such as mammalian cells. Ultimately, this signaling outcome modeling approach can be useful for better understanding of transition from physiological to pathological conditions such as inflammation, various cancers and autoimmune diseases.

**Brief Summary:** Cells are supposed to make correct decisions, i.e., respond properly to various signals and initiate certain cellular functions, based on the signals they receive from the surrounding environment. Due to signal transduction noise, signaling malfunctions or other factors, cells may respond differently to the same input signals, which may result in incorrect cell decisions. Modeling and quantification of decision making processes and signaling outcomes in cells have emerged as important research areas in recent years. Here we present univariate and multivariate data-driven statistical models and methods for analyzing dynamic decision making processes and signaling outcomes. Furthermore, we exemplify the methods using single cell data generated by a p53 system, in wild-type and abnormal cells.

## Introduction

Understanding how cells make decisions in response to input signals is an important challenge in molecular and cell biology. Depending on the signals they receive, cells can adopt different fates. Emergence of single cell data and methods [1-3] has made it possible to study and model the behavior of each cell individually. An important factor that affects cell decisions is biological noise in various organisms [4], which can cause cells to exhibit different behaviors, when receiving the same input signal. For example, under the same stimuli, some cells may decide to survive, whereas others may undergo apoptosis. Signaling outcomes can be affected by genetic and epigenetic regulation and misregulation, leading to errors in signaling outcomes and ensuing cell decisions.

Given the probabilistic nature of cellular decisions [1, 3], it is of interest to have a unified set of statistical metrics and methods to systematically study and characterize the signaling outcomes that may inform them, and determine probabilities associated with different outcomes. Using statistical signal processing and decision theory concepts, recently a framework was introduced by Habibi *et al.* [1], to compute optimal decision thresholds and probabilities for incorrect cell decisions using single cell data. More specifically, in the transcription factor Nuclear Factor κB (NF-κB) pathway regulated by the tumor necrosis factor (TNF) [3], the optimal decision threshold which minimized the decision probability to distinguish between two different TNF levels was computed from data [1]. Probabilities of incorrect cell decisions were computed from data as well.

One goal of this paper is to show how the statistical decision theoretic framework [1] can be used to study other molecular systems and signaling outcomes. The other goal is to extend the decision modeling framework such that one can model and analyze multi-dimensional signaling outcome processes using multi-time point measurements. This allows to incorporate signaling dynamics into decision making analysis. Application of receiver operating characteristic curve as a graphical tool to visualize decisions and outcomes under normal and abnormal conditions is introduced here as well. In this paper, we use the tumor suppressor p53 system, as an example, to present the concepts, metrics and algorithms related to decision making and outcome analysis.

The tumor suppressor p53 is an important transcription factor that is responsible for DNA repair, cell cycle suppression, cell growth control and initiation of apoptosis [5-8]. When a healthy cell is exposed to ionizing radiation (IR), DNA damage occurs [9]. Due to the DNA damage, p53 becomes activated [5, 10, 11] and the cell takes one of two possible actions: it can either survive by repairing the DNA or it can trigger apoptosis [8, 12, 13]. Our focus here is to demonstrate how such outcomes can be systematically modeled. We accomplish this by introducing metrics and methods to evaluate success and failure rates of the signaling outcomes and actions, in response to the DNA damage caused by different IR doses and under various conditions. In order to do this, we collected data using the simulator of Hat *et al.* [9], to obtain single cell data of healthy cells, when different IR doses are applied. Moreover, we collected single cell data of abnormal cells exposed to different IR doses, to measure how the decision making is affected when there is an anomaly in the system, in addition to the DNA damage.

The rest of the paper is organized as follows. First, we briefly explain the p53 system and its response to DNA damage. Then we present decision making and outcome analysis as a hypothesis testing problem on the IR level, define probabilities associated with various decisions, introduce the optimal decision maker, and describe the single cell data used to determine the decision probabilities in the presence of noise and under normal and abnormal conditions. Methods for computing optimal decision thresholds and the associated decision error rates are presented afterwards, using either single, double or multiple time point measurements in individual cells. The latter is particularly useful to understand the effect of temporal variations and dynamical changes. Additionally, receiver operating characteristic curves are computed and presented as useful tools to visualize the tradeoff between decision error rates and how they are affected by decision thresholds and other factors. A comparison between binary and ternary decision making and outcome analysis and their error rates is provided as well. The paper concludes with a summary of the highlights of the methods and their biological implications for understanding signaling outcomes and decisions in the exemplary p53 system, and extensions to other systems.

### Signaling outcomes and decisions in the p53 system when DNA damage occurs: A case study

The transcription factor p53 has a significant role in DNA repair, cell cycle suppression, regulation of cell growth, and initiation of apoptosis [5-8]. It becomes active in response to DNA damage that may occur when the cell is exposed to ionizing radiation (IR), ultraviolet (UV) radiation, heat shock, etc. [5, 10, 11]. In particular, exposure to IR results in DNA double strand breaks (DSBs), which are the most serious DNA lesion. When DSB is not repaired, it can cause cell death or DNA mutations which can propagate to new cell generations [12, 14, 15]. When DNA damage occurs, p53 can assume two phosphorylation states: p53_Arrester_ and p53_Killer_. Afterwards, the p53 system can take two actions: it either suppresses cell cycle until DNA is repaired, if the damage is low and repair is possible; or it can trigger apoptosis if the damage is high and repair is not possible [8, 12, 13]. Herein, we intend to compute decision thresholds and incorrect decision rates when the DNA damages caused by various IR doses occur in a cell. With this goal in mind, we conduct stochastic simulations of cells exposed to different IR doses [9], to obtain in silico single cell data.

Consider the p53 system model [9] shown in Fig 1. The p53 system is activated due to a DNA damage induced by IR. Initially the protein kinase ataxia-telangiectasia mutated (ATM) is activated by the DNA damage [16, 17]. The active ATM phosphorylates Mdm2, which is a p53 inhibitor [18]. The ATM also activates p53 by phosphorylating it to one of its active phosphoforms: p53_Arrester_ which further phosphorylates p53 to the p53_Killer_ form [19-21]. Moreover, the p53_Arrester_ activates the Mdm2 [22] and wild-type p53-induced phosphatase 1 (Wip1) [23, 24]. The active Wip1 inhibits the ATM [25] and dephosphorylates the p53_Killer_ to the p53_Arrester_ form [26]. The p53_Killer_ regulates another phosphatase, phosphatase and tensin homolog (PTEN), which initiates a slow positive feedback loop stabilizing the level of p53 [27]. If DNA damage is large and its repair takes longer time, PTEN accumulates to high levels and inhibits AKT, which may no longer phosphorylate Mdm2. Unphosphorylated Mdm2 remains in cytoplasm and may not target nuclear p53 for degradation. Thus, accumulation of PTEN results in disconnection of negative feedback loop between p53 and Mdm2. The slow positive feedback loop acts as a clock giving cells time to repair DNA, and initiating apoptosis if DNA repair takes too long. The apoptotic module, where transcription of pro-apoptotic proteins is induced, is controlled by p53_Killer_ and Akt that suppresses the apoptosis. When Akt is inhibited by increased level of PTEN, it will no longer suppress the apoptotic module. Thus, the p53_Killer_ will initiate activation of cysteine-aspartic proteases (Caspases), enzymes having essential role in cell death (Fig 1). Since we are interested in the analysis of the signaling outcomes which affect whether the cell survives or triggers apoptosis, we do not consider the cell cycle arrest module (regulated by p53_Arrester_), and focus on the apoptotic module. Simulation files can be found in Hat *et al.* [9] and more detailed information about the p53 system and each component and interaction there can be found in Hat *et al*. [9] and Bogdal *et al.* [29]. More specifically, interested readers can refer to the Supporting Information S1 Text of [9], which includes a summary of mathematical models of the p53 system, a detailed description of the model, a notation guide, and lists of parameters and reactions.

**Fig 1:**
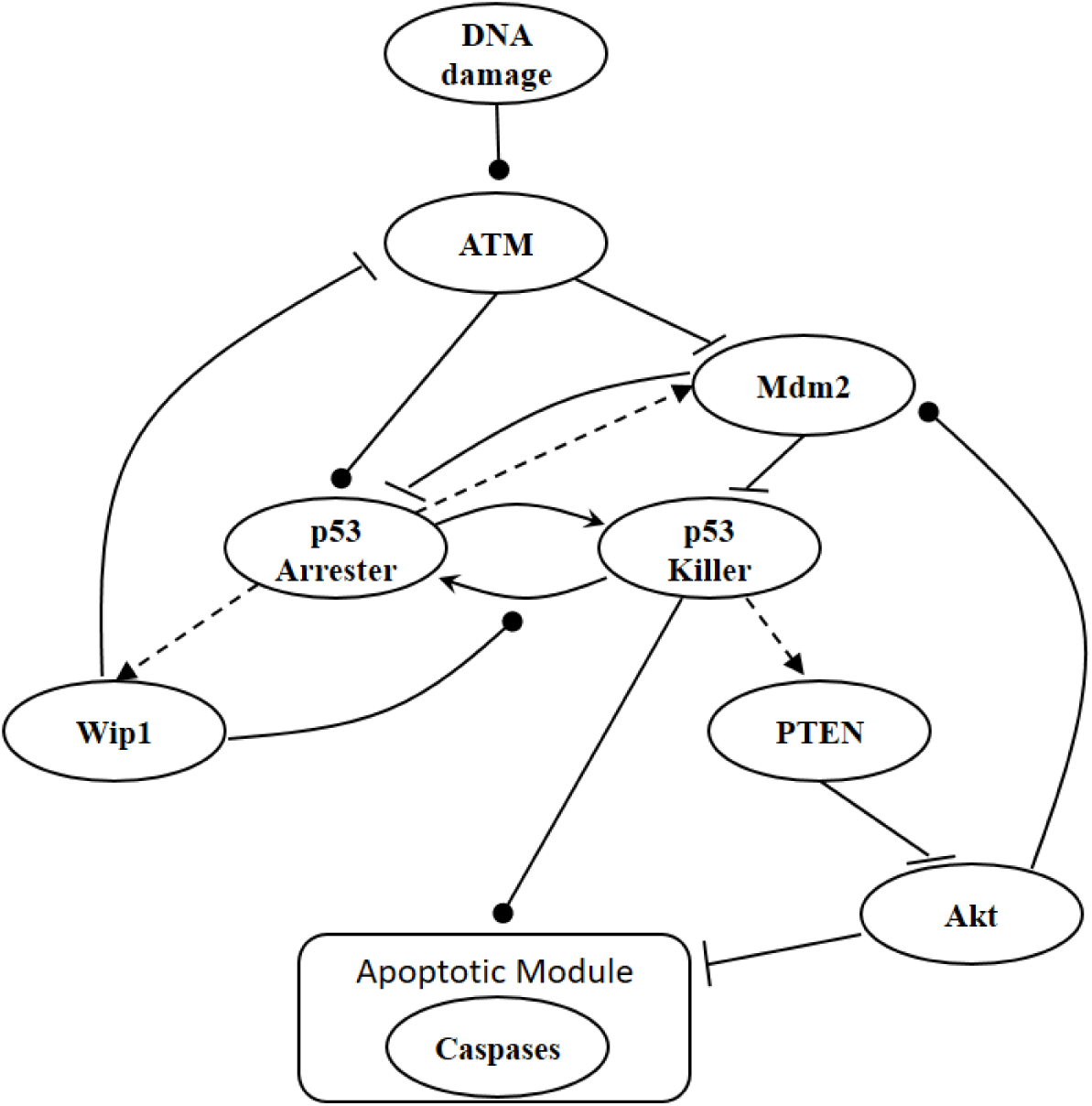
A p53 system model [9]. Arrow-headed dashed lines represent positive transcriptional regulations, arrow-headed solid lines stand for protein transformations, circle-headed solid lines are activatory regulations, and hammer-headed solid lines represent inhibitory regulations. All the molecules and the interactions between them are described in the main body of the paper.

### Decision making and outcome analysis: Hypothesis testing on input signals and optimal decisions with minimum errors

When cells are exposed to radiation, each cell may respond differently due to noise or some other factors. One may decide to survive, whereas another may trigger apoptosis, both under the same IR dose. Given the probabilistic nature of such decisions [1], we can formulate p53-based decision making as a binary hypothesis testing problem, where the decision making system is going to test which of the following two hypotheses is true regarding the applied IR dose, to trigger an action accordingly:

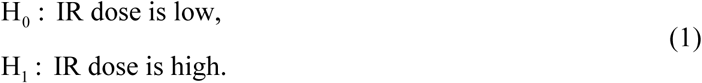

Binary hypothesis testing is observed in other systems, e.g., the TNF/NF-κB system [1].

In response to an IR dose, two types of incorrect decisions can be made. One is deciding that the input IR level is high, whereas in fact it is low (deciding H_1_ when H_0_ is true), which may falsely trigger apoptosis. The other one is deciding that the input IR level is low, whereas in fact it is high (deciding H_0_ when H_1_ is true), which may result in missing apoptosis. These two erroneous decisions can be called as *false alarm* and *miss event*, respectively, and their probabilities can be defined as:

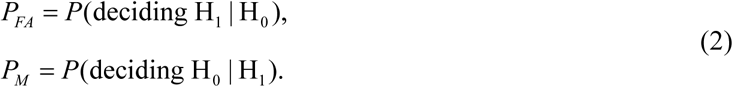

The overall error probability *P*_*E*_ of making decisions is a combination of *P*_*FA*_ and *P*_*M*_ :

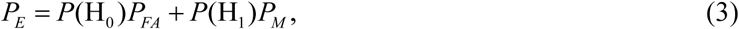

Where *P*(H_0_) and *P*(H_1_) are prior probabilities of H_0_ and H_1_, respectively. Note that as mentioned in the Introduction section, IR causes DNA damage. Therefore, one can instead formulate the p53-based decision making process as a binary hypothesis testing on DNA damage being low or high, and define the associated false alarm and miss events probabilities accordingly.

The optimal decision making system which minimizes the above *P*_*E*_ is the one that compares probabilities of observed data under the hypotheses H_0_ and H_1_ [30]. More precisely, suppose that *x* is the observation and *p*(*x* | H_0_) and *p*(*x* | H_1_) are the conditional probability density functions (PDFs) of *x* under H_0_ and H_1_, respectively. Also consider equi-probable hypotheses, i.e., *P*(H_0_) = *P*(H_1_) = 0.5, which is a reasonable assumption in the absence of prior information on the possibilities of H_0_ and H_1_. Then, the optimal system decides H_1_ if *p*(*x* | H_1_) > *p*(*x* | H_0_), otherwise, it decides H_0_. This means that the hypothesis with the highest likelihood is decided. This decision is called the maximum likelihood decision [30].

### Single cell data of the p53 system exposed to ionizing radiation

To calculate the error probabilities in Equation (2), we use PTEN level as the decision variable because when unrepairable DNA damage occurs, the activated p53 triggers pro-apoptotic phosphatase PTEN [27], and PTEN initiates apoptosis [28]. It has also been shown by Hat *et al.* [9] that PTEN is a decent predictor of cell fate. After specifying the decision variable, we use the stochastic simulator of Hat *et al.* [9] to generate 5000 cells for each IR dose. The stochastic simulation has three phases. The first phase is the “equilibrium phase” where we simulate 2 weeks of cell behavior when no IR dose is applied. The second phase is called “irradiation phase” in which 10 minutes of IR dose is applied. The last phase is called “relaxation phase” in which we simulate 72 hours of cell behavior after it is exposed to 10 minutes of IR. When IR dose increases, apoptotic cell percentage increases as well [9] (Fig 2). For more details on the simulation phases, see supporting files of Hat *et al.* [9]. In order to decide whether a cell is apoptotic or not, we check active caspase level in 72 hours after the irradiation phase, and compare it with the threshold of 0.5×10^5^ suggested in Hat *et al.* [9]. Cells with the level of active caspase higher (or lower) than the threshold of 0.5×10^5^ are considered to be apoptotic (or surviving).

**Fig 2:**
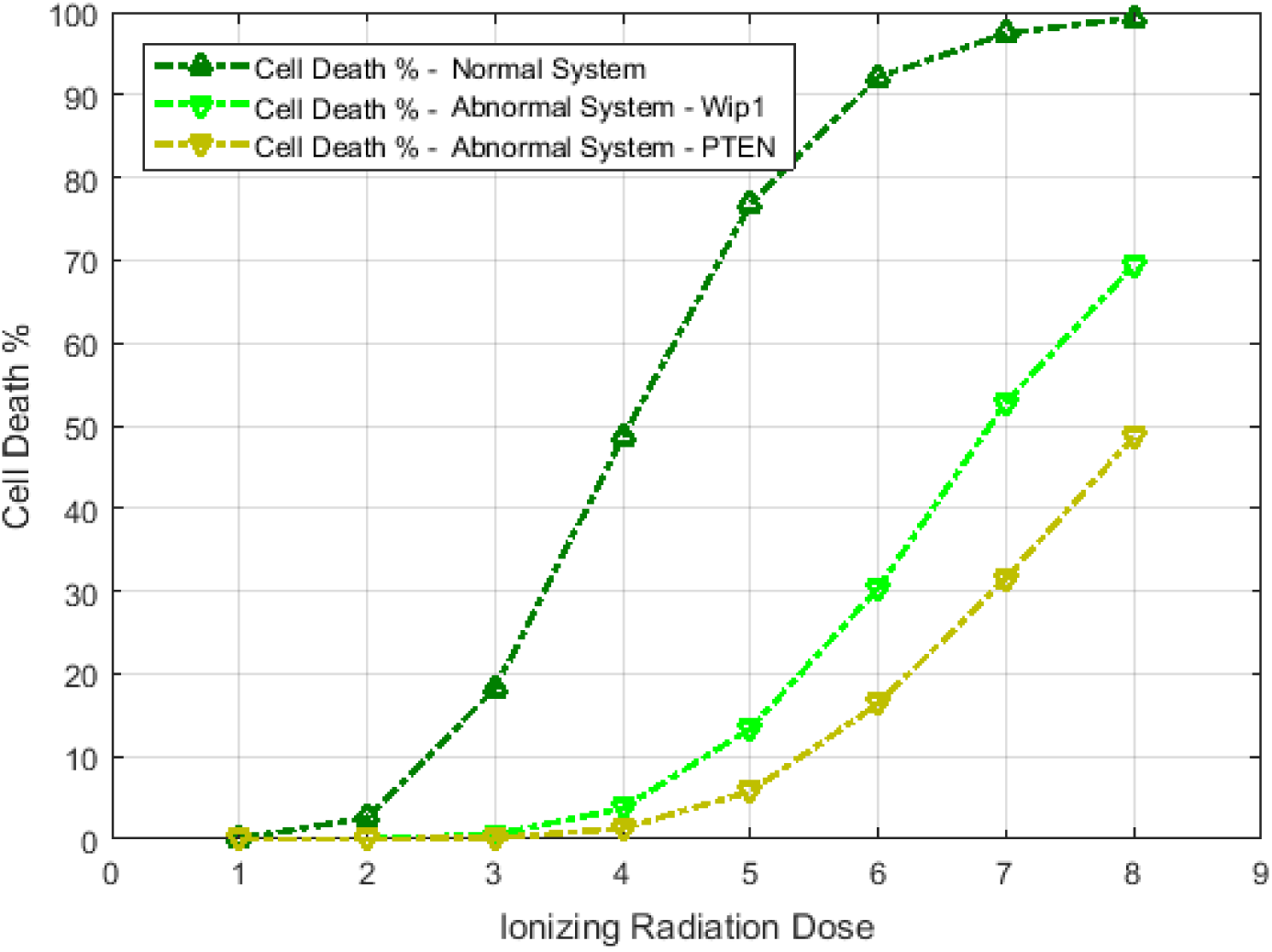
Cell death percentage versus ionizing radiation (IR) dose in both normal and abnormal p53 systems. The dark green curve at the top represents a normal p53 system with no perturbation, whereas the other two curves correspond to p53 systems behaving abnormally due to Wip1 or PTEN perturbations.

The data of normal cells includes eight sets of PTEN levels in 5000 cells, which correspond to eight doses of IR = 1, 2, 3, 4, 5, 6, 7 and 8 Gy. Here Gy stands for Gray, the unit of radiation dose, and 1 Gy is 1 Joule of energy absorbed by 1 kg of tissue. We focus our analysis on low IR versus high IR hypothesis testing, to see how accurately it can be decided whether the applied radiation level is low or high. We consider IR = 1 Gy as the low dose, whereas the higher dose can be IR = 2, 3, 4, 5, 6, 7 or 8 Gy. More specifically, scenarios in which signaling outcomes are analyzed are 1 vs. 2 Gy, 1 vs. 3 Gy, 1 vs. 4 Gy, 1 vs. 5 Gy, 1 vs. 6 Gy, 1 vs. 7 Gy, and 1 vs. 8 Gy. We quantitatively study in which of these scenarios more erroneous decisions are made. We also determine to what extent decision between responses to low and high IR levels depends on the input IR separation. We conduct these studies by computing the optimal decision threshold in each scenario using the PTEN data, following the maximum likelihood principle that provides the best decisions, i.e., smallest decision error probabilities. We also compute numerical values of the decision error probabilities using the PTEN data.

In addition to the analysis of erroneous decision making and incorrect signaling outcomes in normal cells mentioned above, we analyze them in abnormal cells as well, where there is a dysfunctional molecule in the p53 system. Wip1 is one of the key regulatory pro-survival phosphatases [23] in the p53 system (Fig 1). If the DNA damage can be repaired, then Wip1 expression returns the cell to the pre-stress state from cell-cycle arrest [23, 32]. It has been observed that elevated Wip1 level exists in multiple human cancer types such as breast, lung, pancreas, bladder, and liver cancer [33-38]. Therefore, to obtain abnormal cells, we generate cells with increased Wip1 synthesis rate. In normal cells, Wip1 synthesis rate is about 0.1 [9], and here we increase it to 0.15, a 50% increase, to reproduce abnormality. This increase in the Wip1 synthesis rate causes a significant decrease in the cell death percentage (Fig 2), which can be considered as an abnormal cell state. In addition to Wip1, we analyze abnormal cellular state caused by PTEN abnormalities. It has been observed that attenuated PTEN levels exist in MCF-7, a non-invasive form of human breast cancer cells [39]. Therefore, it is of interest to see how the abnormal PTEN level affects signaling outcomes in the p53 system. To study this, we generate abnormal cells by decreasing PTEN synthesis rate. In healthy cells, the PTEN synthesis rate is about 0.03 [9]. Here we decrease it to 0.015, a 50% decrease, to reproduce abnormality. We observe a considerable decrease in the cell death percentage (Fig 2), representing an abnormal cellular state.

### Univariate analysis: Methods for computing decision thresholds and decision error rates using *single* time point measurements in individual cells

In this section, we analyze PTEN levels of 5000 cells measured in 72 hours after the irradiation phase. It has been observed that PTEN levels of both apoptotic and surviving cells become very distinct in 72 hours after 10 minutes of IR application [9] (decision analysis based on PTEN levels at other time instants, as well as multiple time instants are presented in other sections).

Histograms of natural logarithm, ln, of PTEN levels for IR = 1, 2 Gy data sets and IR = 1, 8 Gy data sets are shown in Fig 3A and Fig 3C, respectively. As presented in Fig 3B and Fig 3D, Gaussian PDFs whose means and variances are estimated from the data, reasonably represent the histograms. This indicates that the PTEN data can be reasonably approximated by lognormal PDF. Due to the mathematical convenience of working with Gaussian PDFs and variables, especially for multivariate analysis of multiple time point data discussed later, we continue working with the logarithm of the PTEN data. Let *x* = ln(PTEN) be the Gaussian variable of interest with mean *μ* and variance *σ*^2^, i.e., *x* ∼ ***𝒩*** (*μ,σ*^2^) where ***𝒩*** stands for the following normal or Gaussian PDF:

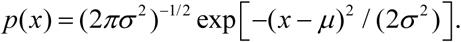

**Fig 3:**
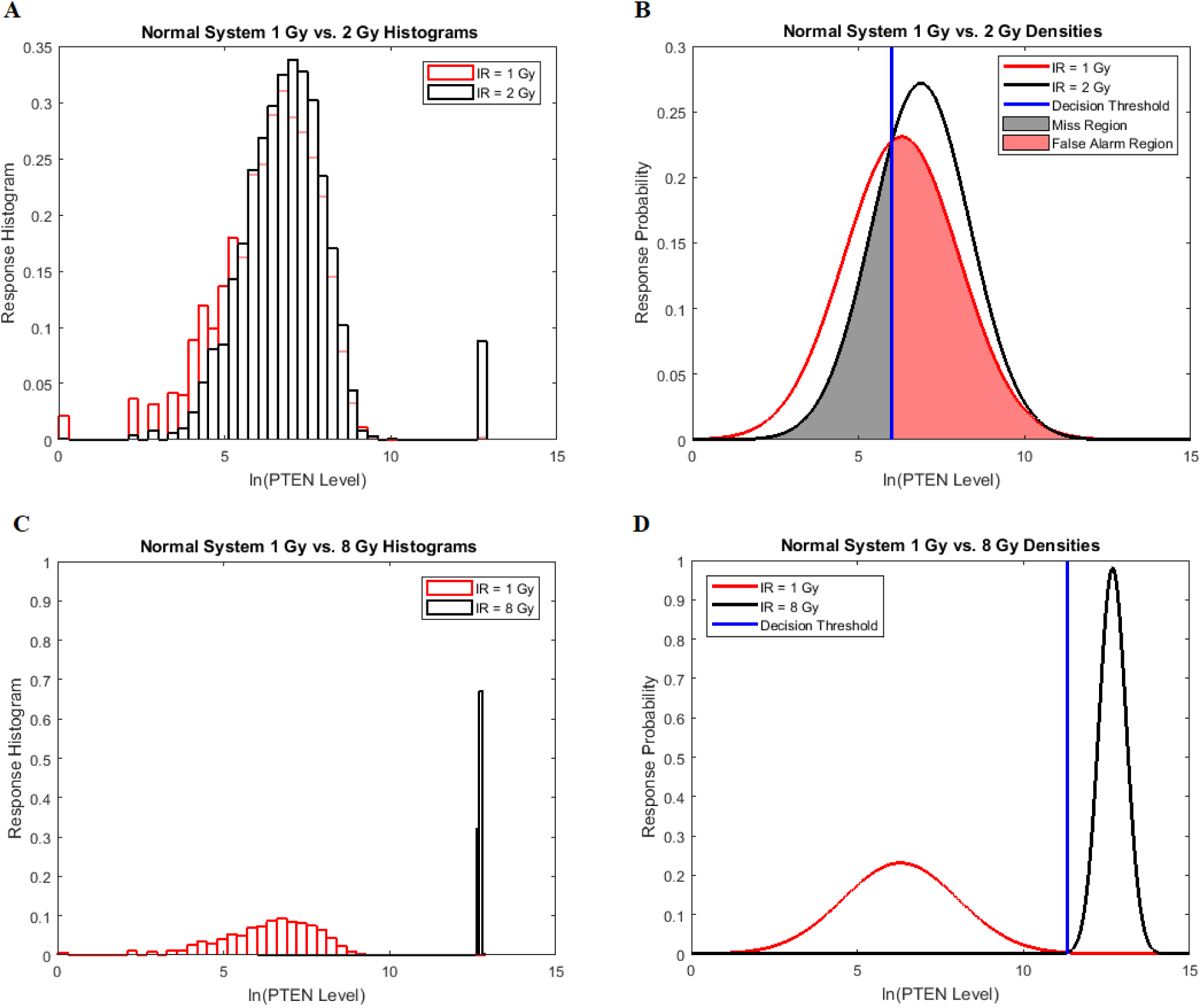
Univariate decision making and signaling outcome analysis in the normal p53 system based on PTEN response distributions. **(A)** Histograms of PTEN levels of cells under IR = 1 Gy and IR = 2 Gy doses. **(B)** Gaussian probability density functions (PDFs) for PTEN levels of cells under IR = 1 Gy and IR = 2 Gy doses, together with the optimal maximum likelihood decision threshold which minimizes the total decision error probability. **(C)** Histograms of PTEN levels of cells under IR = 1 Gy and IR = 8 Gy doses. **(D)** Gaussian PDFs for PTEN levels of cells under IR = 1 Gy and IR = 8 Gy doses, together with the optimal maximum likelihood decision threshold which minimizes the total decision error probability.

The Gaussian PDFs shown in Fig 3 are indeed the conditional PDFs *p*(*x* | H_0_) and *p*(*x* | H_1_) under the hypotheses H_0_ and H_1_ defined earlier in Equation (1). For example, in Fig 3B, H_0_ and H_1_ correspond to IR = 1 Gy and IR = 2 Gy doses, respectively, and the red and black curves in there are the conditional PDFs *p*(*x* | H_0_) and *p*(*x* | H_1_), respectively.

#### The optimal maximum likelihood decision making system

Recall our two hypotheses previously defined in (1). The optimal decision maker, which minimizes the overall error probability *P*_*E*_ in (3), compares the conditional likelihood ratio *L*(*x*) = *p*(*x* | H_1_) / *p*(*x* | H_0_) with the ratio *γ* = *P*(H_0_) / *P*(H_1_) [1]. The system decides H_1_ if *L*(*x*) > *γ*. If the hypotheses are equi-probable, i.e., *P*(H_0_) = *P*(H_1_) = 0.5, then the optimal system decides H_1_ if *p*(*x* | H_1_) > *p*(*x* | H_0_).

#### The optimal decision threshold

To find the optimal decision threshold, we need to solve the equation *L*(*x*) = *γ*, i.e., *P*(H_1_) *p*(*x* | H_1_) = *P*(H_0_) *p*(*x* | H_0_), for *x*. When H_0_ and H_1_ are equi-probable, the threshold equation to be solved simplifies to *L*(*x*) = 1, i.e., *p*(*x* | H_1_) = *p*(*x* | H_0_).

#### The decision error probabilities

Once the optimal decision threshold is determined, it can be used to compute false alarm and miss decision error probabilities, by integrating the conditional PDFs of data over error regions. More specifically, using the conditional PDFs *p*(*x* | H_0_) and *p*(*x* | H_1_) representing the response probabilities of the ln of PTEN levels under the two hypotheses, Equation (2) can be written as [1]:

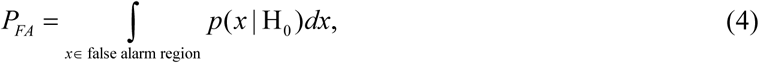

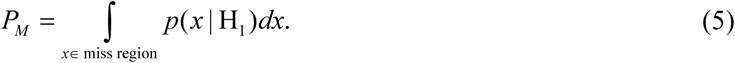

The false alarm region in (4) is defined by {*x* : *p*(*x* | H_1_) > *p*(*x* | H_0_)} when H_0_ is true, whereas the miss region in (5) is defined by {*x* : *p*(*x* | H_0_) > *p*(*x* | H_1_)} when H_1_ is true. By substituting *PFA* and *P*_*M*_ in Equation (3) the overall error probability *P*_*E*_ can be obtained.

#### The Gaussian data model to compute the optimal decision threshold

Here we focus on Fig 3B as an example, where two Gaussian PDFs are shown for *x* = ln(PTEN), the natural logarithm of PTEN levels in the two data sets of IR = 1 Gy and IR = 2 Gy, with each data set consisting of 5000 cells. Let 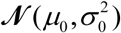 and 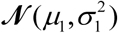 represent the Gaussian PDFs that correspond to the IR = 1 Gy and IR = 2 Gy data sets, respectively, where 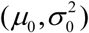 and 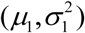 are mean/variance pairs estimated from their associated data sets. The optimal maximum likelihood decision threshold in Fig 3B is at the intersection of the two PDFs, and can be computed by solving the equation *p*(*x* | H_0_) = *p*(*x* | H_1_) written below:

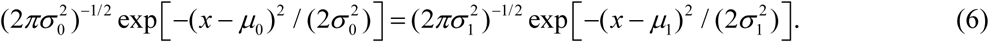

By multiplying both sides by 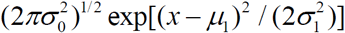 and then taking natural logarithm of both sides, (6) can be written in the following quadratic equation form [1]:

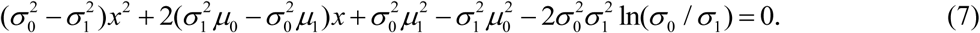

Equation (7) is derived assuming our hypotheses are equi-probable, i.e., *P*(H_0_) = *P*(H_1_) = 0.5, as mentioned before. The solution of Equation (7) gives the optimal decision threshold *PTEN*_th_, located at the intersection of the two PDFs for IR = 1 Gy and IR = 2 Gy doses in Fig 3B (the italic style is adopted to clarify that the threshold is related to the logarithm of PTEN data). Interestingly, for equal variances, solution of Equation (7) for the optimal decision threshold simplifies to the average of the means, i.e., (*μ*_0_ + *μ*_1_) / 2, which intuitively makes sense. For other prior probabilities and PDF models, the optimal threshold can be obtained similarly, by solving the equation *P*(H_0_) *p*(*x* | H_0_) = *P*(H_1_) *p*(*x* | H_1_) for *x*.

#### The Gaussian data model to compute the decision error probabilities

Using the *PTEN*_th_ obtained by solving Equation (7) and using the Gaussian PDFs, Equations (4) and (5) for the false alarm and miss error probabilities can be written as:

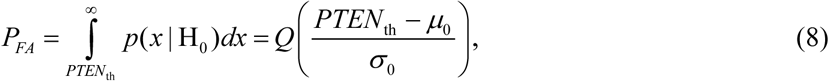

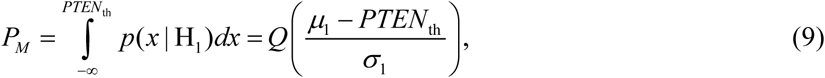

where *Q*(*η*) is tail probability of the standard Gaussian PDF ***𝒩*** (0,1) :

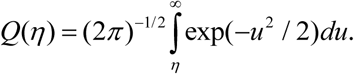

Equation (8) represents area of the pink region in Fig 3B under the tail of the IR = 1 Gy PDF, beyond the *PTEN*_th_ threshold. In this region of *x* > *PTEN*_th_ we have *p*(*x* | H_1_) > *p*(*x* | H_0_), while H_0_ is true. This is the *false alarm* region for which we have computed *P*_*FA*_ = 0.57 in Fig 3B. On the other hand, Equation (9) represents area of the gray region in Fig 3B under the tail of the IR = 2 Gy PDF, below the *PTEN*_th_ threshold. In this region of *x* < *PTEN*_th_ we have *p*(*x* | H_0_) > *p*(*x* | H_1_), while H_1_ is true. This is the *miss* region for which we have computed *P*_*M*_ = 0.28 in Fig 3B. After computing *P*_*FA*_ and *P*_*M*_, we can now compute the overall error probability *P*_*E*_ using Equation (3), which results in *P*_*E*_ = (*P*_*FA*_ + *P*_*M*_) / 2 = 0.43. Similarly, by computing Equations (8) and (9) for the 1 vs. 8 Gy scenario we obtain *P*_*E*_ = 0.001 (Fig 3D). Based on the results of 1 vs. 2 Gy and 1 vs. 8 Gy decision scenarios, it can be concluded that when the difference between the two applied IR doses increases, the overall decision error probability *P*_*E*_ decreases. This is mainly because the two response PDFs become more distinct with less overlap, as the difference between the two applied IR doses increases.

For some cases such as 1 vs. 3, 4, 5 and 6 Gy IR doses, some data sets need to be modeled by a mixture of Gaussian PDFs due to bistable behavior of p53 system and hence cells’ bimodal histograms. Still the same underlying theory and proposed framework hold. Nevertheless, in what follows we explain how to determine the optimal decision thresholds and how to compute the decision error probabilities when using a mixture model, for the 1 vs. 4 Gy scenario.

Histograms of natural logarithm of PTEN levels for IR = 1, 4 Gy data sets are shown in Fig 4A. We notice that while 1 Gy data histogram is unimodal, histogram of 4 Gy data is bimodal. Therefore, for the 1 Gy data we use a single Gaussian PDF as before, whereas for the 4 Gy data we utilize a mixture of two Gaussian PDFs. More specifically, we consider 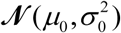 for H to represent the single Gaussian PDF that corresponds to the IR = 1 Gy data, whereas we use 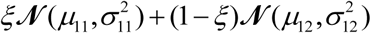 for H_1_, with 0 ≤ *ξ* ≤1 being the mixing parameter, to represent the mixture of two Gaussian PDFs which correspond to the IR = 4 Gy data set. The mean and variance 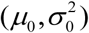 are estimated from the 1 Gy data and the associated single Gaussian PDF is shown in Fig 4B. Furthermore, the means and variances 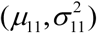 and 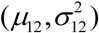 and the mixing parameter *ξ* are estimated from the 4 Gy data using the MATLAB command “fitgmdist” which implements the iterative Expectation-Maximization (EM) algorithm. The resulting mixture of two Gaussian PDFs is shown in Fig 4B.

**Fig 4:**
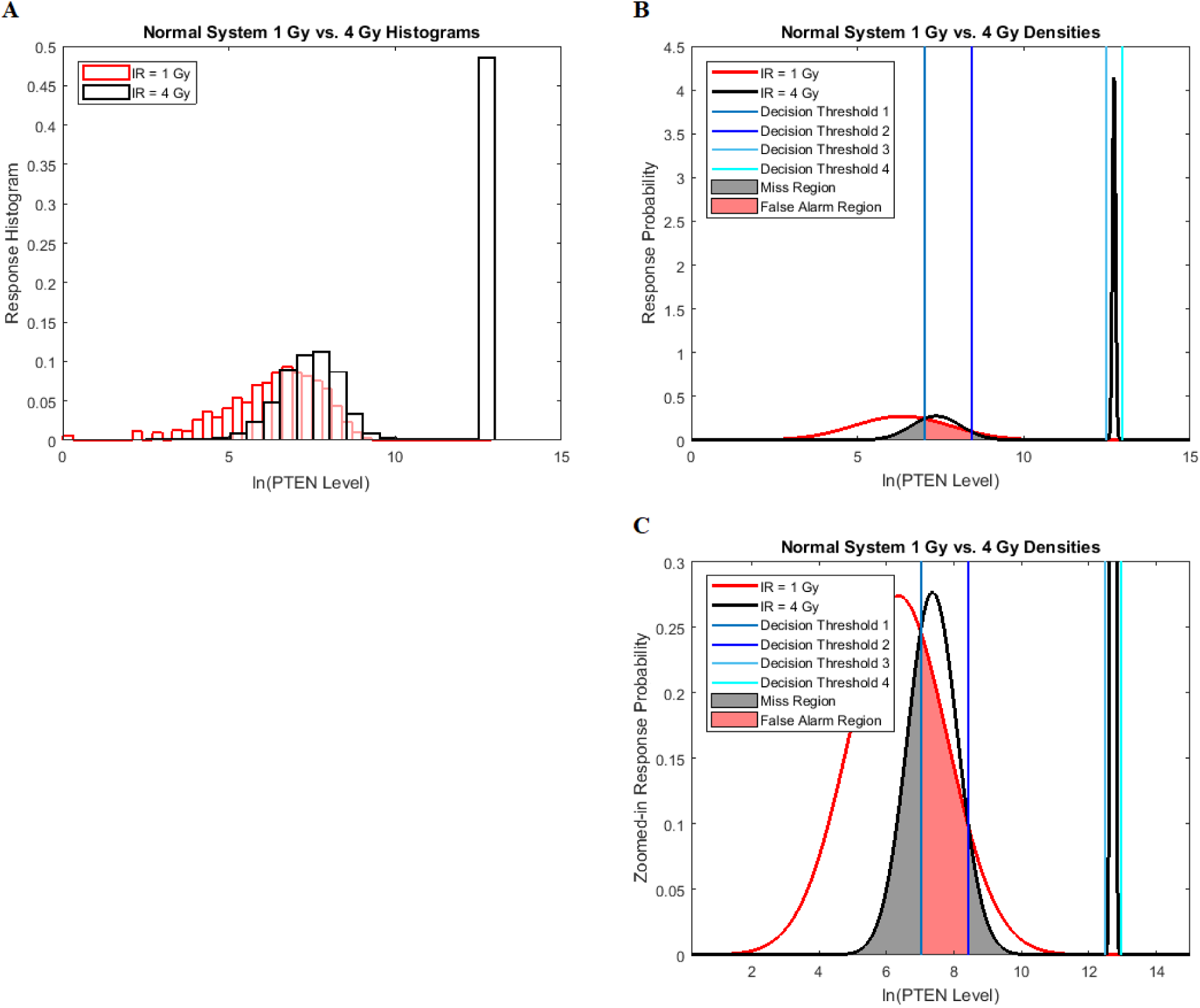
Univariate decision making and signaling outcome analysis in the normal p53 system when a PTEN response distribution is bimodal. **(A)** Histograms of PTEN levels of cells under IR = 1 Gy and IR = 4 Gy doses. **(B)** A Gaussian probability density function (PDF) for PTEN levels of cells under IR = 1 Gy and a mixture of two Gaussian PDFs for PTEN levels of cells under IR = 4 Gy doses, together with the optimal maximum likelihood decision thresholds which minimize the total decision error probability. **(C)** Zoomed-in view of panel B.

Similar to the previous scenarios, the optimal maximum likelihood decision thresholds shown in Fig 4B for equi-probable hypotheses are at the intersections of the conditional PDFs *p*(*x* | H_0_) and *p*(*x* | H_1_), the latter being a Gaussian mixture for the 4 Gy data. Note that here solving the equation *p*(*x* | H_0_) = *p*(*x* | H_1_) results in four solutions for *x*, that is why there are four decision thresholds, *PTEN*_thi_, i = 1, 2,3, 4 in Fig 4B (Note that each decision threshold *PTEN*_thi_ is listed as “Decision Threshold i” in Fig 4).

To compute the decision error probabilities, the false alarm and miss probabilities *P*_*FA*_ and *P*_*M*_ need to be calculated using Equation (4) and Equation (5), respectively. Since there are four decision thresholds in this case, integration has to be performed over multiple regions, which results in lengthy expressions. However, note that as can be seen in Fig 4B and its zoomed-in view in Fig 4C, PDFs for low dose (red) and the lower Gaussian mode for the high dose (black) assume very small values as they reach the third threshold. Therefore, their contributions to possible error events around the third and fourth thresholds are negligible (later this is shown numerically). Similarly, given the very small variance of the higher Gaussian mode of the PDF for the high dose, this PDF is substantially different from zero only between the third and fourth thresholds. Consequently, the contribution of the PDF of this mode to possible errors around the third and fourth thresholds is negligible as well. Overall, as just explained, optimal decision when *PTEN*_th3_ < *x* < *PTEN*_th4_ is H_1_ with no decision error, whereas for *x* < *PTEN*_th1_, *PTEN*_th1_ < *x* < *PTEN*_th2_ and *PTEN*_th2_ < *x* < *PTEN*_th3_, optimal decisions are H_0_, H_1_ and H_0_, respectively, with the following decision error probabilities:

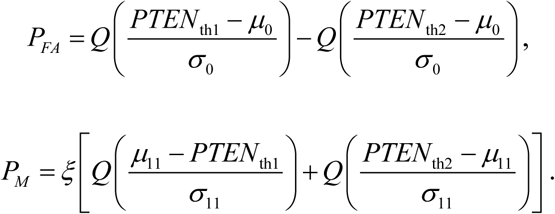

The *P*_*FA*_ expression corresponds to the pink region in Fig 4C, whereas the two *Q* functions in the *P*_*M*_ expression correspond to the two gray regions in Fig 4C, respectively. Using the data, computed numerical values are *ξ* = 0.51, *P*_*FA*_ = 0.28 − 0.06 ≈ 0.22, *P*_*M*_ = 0.51[0.41 + 0.07] ≈ 0.25 and *P*_*E*_ ≈ 0.24, the last one being calculated using Equation (3).

As an example of a negligible decision error probability around the third and fourth thresholds mentioned earlier, consider the area under the red Gaussian PDF *p*(*x* | H_0_) in Fig 4C for *PTEN*_th3_ < *x* < *PTEN*_th4_. While not visible due to being very small, it can be understood that the aforementioned area is a false alarm probability of deciding H_1_, although H_0_ is true. Numerical value of this false alarm probability is *Q* ((*PTEN*_th3_ − *μ*_0_) / *σ*_0_) − *Q* ((*PTEN*_th4_ − *μ*_0_) / *σ*_0_) = 1.2 ×10^−5^ − 2.8 ×10^−6^ ≈ 0, which is negligible compared to *P*_*FA*_ ≈ 0.22 calculated in the previous paragraph.

#### Abnormal p53 systems

To see how an abnormality in the p53 system affects the decision making and signaling outcomes, we calculate *P*_*E*_ values when Wip1 synthesis rate is elevated by 50% from 0.1 to 0.15 (Fig 5), as mentioned previously. As suggested by Habibi *et al.* [1], decision thresholds are modeled to be those of the normal cells. This implies that abnormal cells are not aware of the abnormality, and therefore erroneously use the previous threshold. As we see later, this increases decision error probabilities, a behavior that can be anticipated from abnormal cells. Using Equations (8), (9) and (3), *P*_*FA*_, *P*_*M*_ and *P*_*E*_ are computed: *P*_*E*_ = 0.44 is obtained for 1 vs. 2 Gy scenario (Fig 5A), and *P*_*E*_ = 0.16 is obtained for 1 vs. 8 Gy scenario (Fig 5B). Compared to the normal system results, the overall error probability is significantly higher for the abnormal system under the 1 vs. 8 Gy scenario (we observe that *P*_*E*_ = 0.001 of normal cells markedly increases to *P*_*E*_ = 0.16 in abnormal cells). The reason is that when the Wip1 synthesis rate is increased, the two response PDF curves significantly overlap (notice the overlap between the left-side component of the IR = 8 Gy PDF with the IR = 1 Gy PDF in Fig 5B). This is while in normal cells they had almost no overlap (Fig 3D).

**Fig 5:**
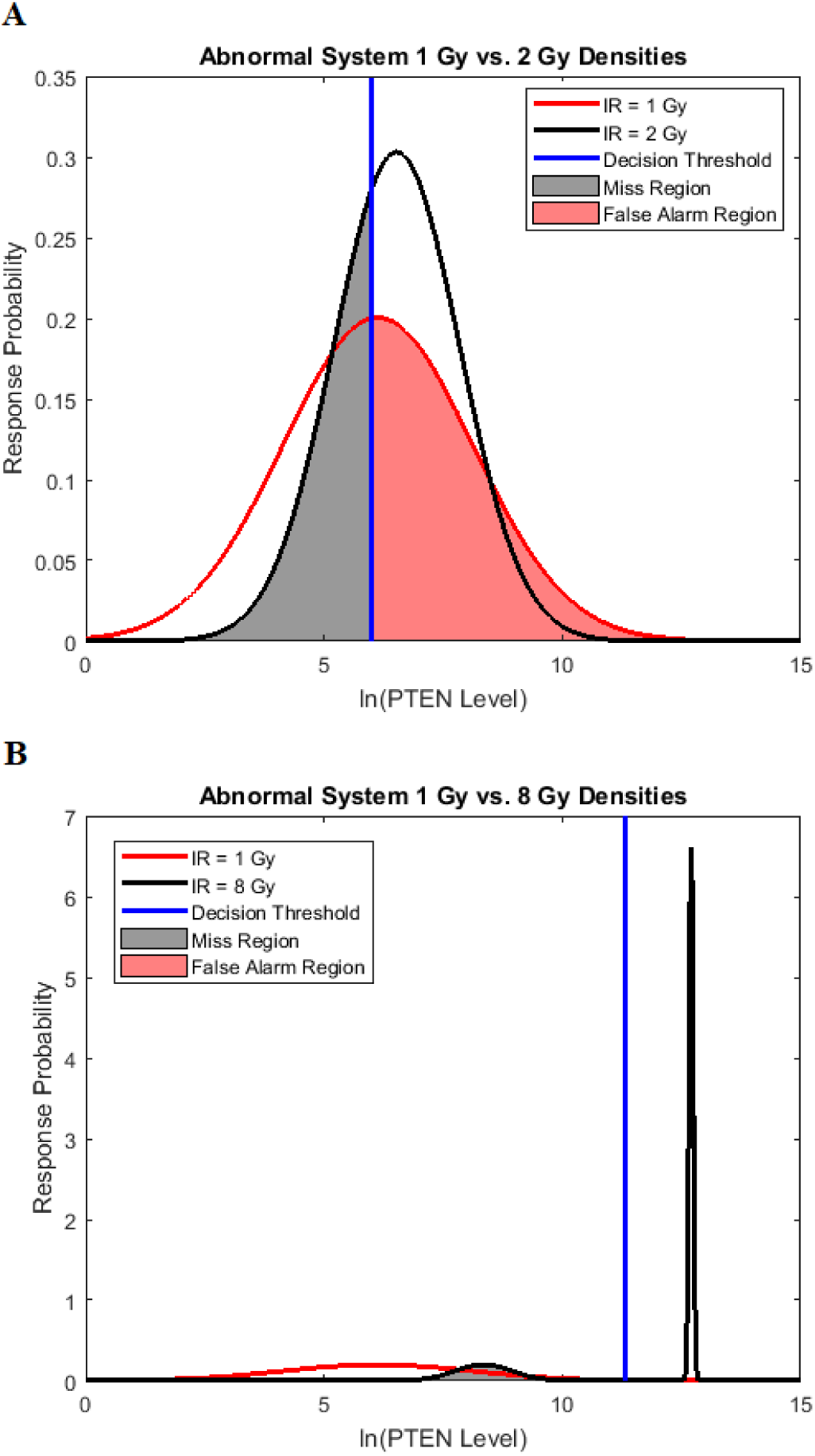
Univariate decision making and signaling outcome analysis in an abnormal p53 system, with increased Wip1 synthesis rate, based on PTEN response distributions. **(A)** Gaussian probability density functions (PDFs) for PTEN levels of abnormal cells under IR = 1 Gy and IR = 2 Gy doses, together with the decision threshold of normal cells. This implies that in abnormal cells the previous decision threshold is erroneously used [1]. As discussed later, this increases decision error probabilities, a behavior that can be anticipated from abnormal cells. **(B)** A Gaussian PDF for PTEN levels of abnormal cells under IR = 1 Gy dose and a mixture of two Gaussian PDFs for PTEN levels of abnormal cells under IR = 8 Gy dose, together with the decision threshold of normal cells.

Similarly, we compute error probabilities for the other abnormal p53 system we mentioned previously, generated by the PTEN synthesis rate reduced from 0.03 to 0.015 (50% reduction). Error probabilities for this abnormality for all different radiation exposure scenarios of 1 vs. 2 Gy up to 1 vs. 8 Gy are shown in Fig 6. For comparison, error probabilities for the Wip1-perturbed abnormal p53 system and also the normal p53 system are provided in Fig 6 as well. We observe that as the difference between the two applied IR doses increases, decision error probability in normal cells drops significantly. This is while in abnormal cells, decision error probabilities remain high. These signaling outcomes might be correlated with the observation that cell death percentages in abnormal systems are considerably lower than the normal system, even when the radiation dose increases (Fig 2). This could indicate that abnormal cells do not respond to IR levels properly and hence, decisions and signaling outcomes affecting apoptosis and survival become more erroneous. Care should be taken that these specific observations are based on the low versus high IR, e.g. d_0_ vs d_1_ IR hypothesis testing formulation where the low IR dose is fixed to 1 Gy (d_0_ = 1 Gy) and the high IR dose is ranging from 2 Gy up to 8 Gy (d_1_ = 2, 3, …, 8 Gy) in the p53 system, that is considered in this paper as an example. These observations may not be generalized to other selections of the low d_0_ and high d_1_ IR doses or other hypothesis testing formulations, case studies or signaling networks. However, the proposed framework and its analytical tools, whose introduction has been the main goal of this paper, can be similarly used.

**Fig 6:**
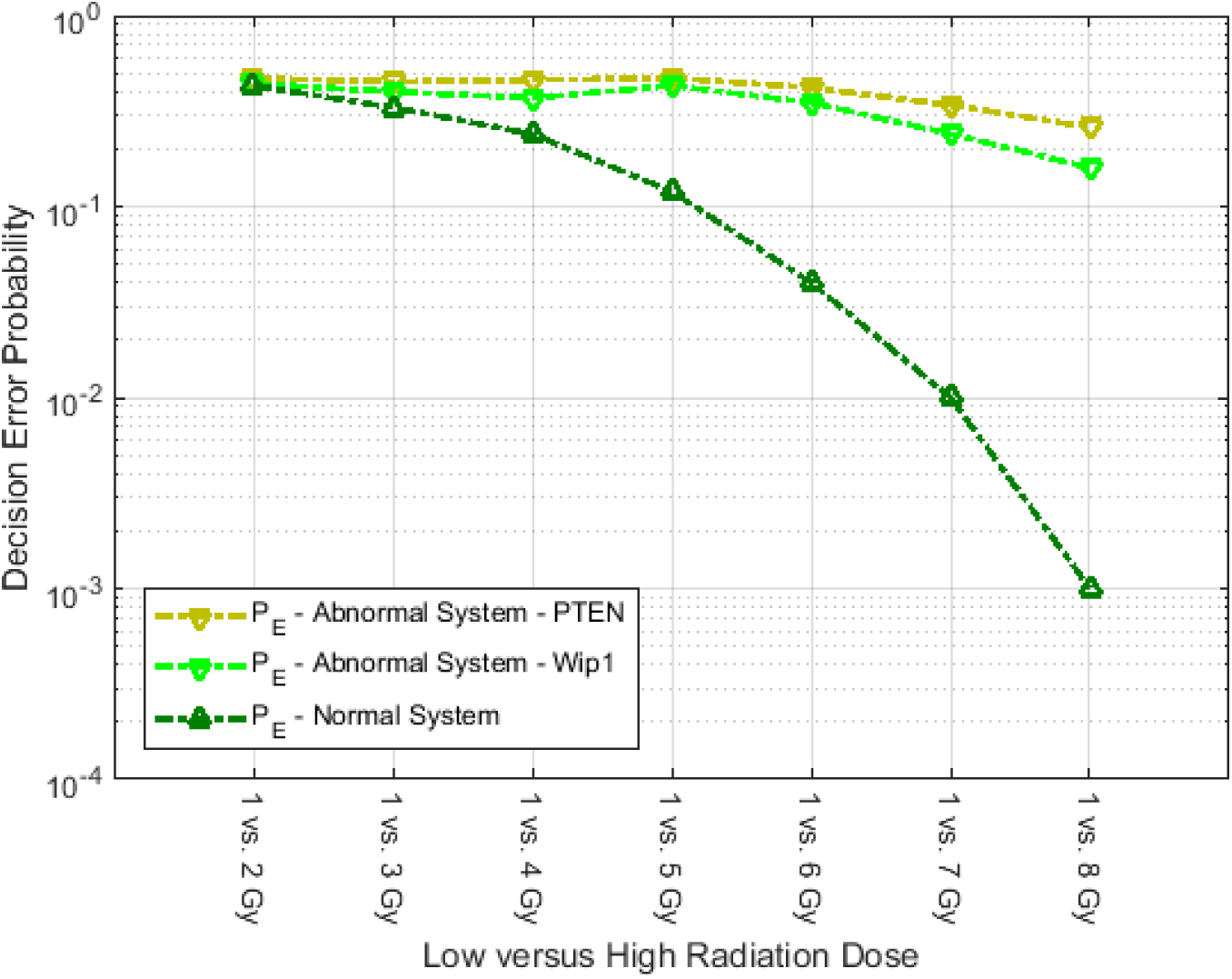
Decision error probabilities for several low IR versus high IR scenarios. The “Abnormal System – PTEN” legend refers to a p53 system whose PTEN synthesis rate is decreased by 50%, compared to its nominal value. The “Abnormal System – Wip1” legend refers to a p53 system whose Wip1 synthesis rate is increased by 50%, compared to its nominal value. Smaller decision error probabilities in the normal system are noteworthy.

### Decision and signaling outcome analysis using receiver operating characteristic (ROC) curves

In this subsection, we show how to analyze performance of a decision maker using receiver operating characteristic (ROC) curves. The ROC curve is developed to visualize the performance of decision making systems [30, 40], and is a graph of probability of detection, *P*_*D*_ = 1 − *P*_*M*_, versus the probability of false alarm, *P*_*FA*_. In Fig 7 we present ROC curves for both the normal p53 system (Fig 7A) and the abnormal p53 system (Fig 7B) whose Wip1 synthesis rate is elevated, for these two low vs. high IR decision making scenarios: 1 vs. 2 Gy and 1 vs. 8 Gy. The theoretical ROC curves in Fig 7 are graphed using the false alarm and miss decision error probability formulas in Equations (8) and (9), respectively, with *μ* s, *σ* s and the thresholds estimated from the data. The empirical ROC curves in Fig 7 are graphed by using the data sets directly, using the MATLAB command “perfcurve”. We observe that the theoretical and empirical ROCs are nearly the same. Therefore, in what follows, we focus on the theoretical ROC curves, to explain concepts and results.

**Fig 7:**
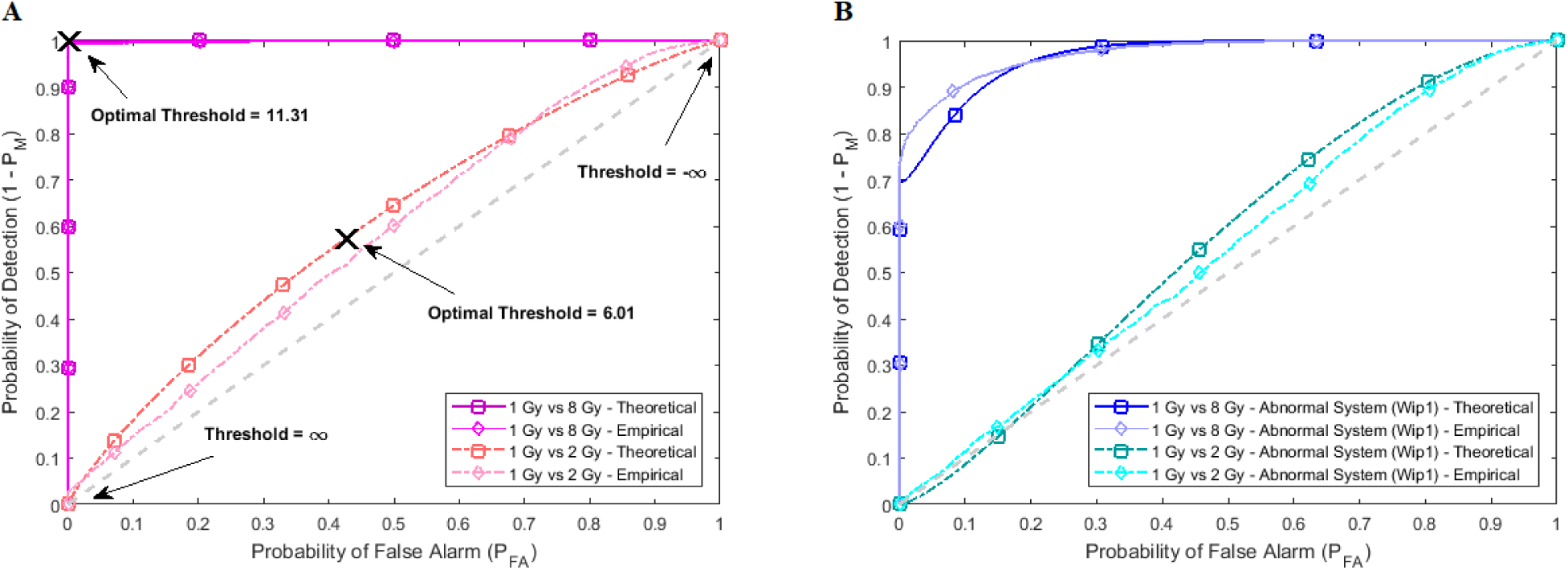
Empirical and theoretical receiver operating characteristic (ROC) curves for both normal and abnormal p53 systems. The theoretical ROC curves labeled by □ are obtained from the Gaussian and mixture of Gaussians data models and formulas whose parameters are estimated from the data, whereas the empirical ROC curves labeled by ◊ are obtained directly from the data. We observe that the theoretical and empirical ROCs are nearly the same. Note that Threshold = ln(PTEN Level) in the figures. **(A)** ROC curves of 1 vs. 2 Gy and 1 vs. 8 Gy radiation scenarios for the normal system. **(B)** ROC curves of 1 vs. 2 Gy and 1 vs. 8 Gy radiation scenarios for the Wip1-perturbed abnormal system.

A ROC curve is above a 45° diagonal line [30], the gray dashed line in Fig 7. In our study it represents the worst possible decision maker, i.e., a decision making system that does not use the data and instead randomly decides if the applied IR dose is low or high, by just flipping a coin. The 45° line is indeed a reference to judge the performance of a decision making system. A ROC curve far away from the 45° reference line indicates a good decision maker. Each point on a ROC curve represents a (*P*_*FA*_, *P*_*D*_) pair that corresponds to a certain decision threshold. Other properties of ROC curves can be found in Van Trees *et al.* [40]. The “**×**” marks in Fig 7A show the optimal (*P*_*FA*_, *P*_*D*_) points that correspond to the optimal decision thresholds shown in Fig 3B and Fig 3D, previously computed using Equation (7) for the 1 vs. 2 Gy and 1 vs. 8 Gy scenarios, respectively.

Based on the normal p53 system ROC curves in Fig 7A, we observe that decisions are made better under the 1 vs. 8 Gy scenario, because of its ROC curve being very far from the 45° reference line, compared to the 1 vs. 2 Gy case whose ROC curve is much closer to the 45° reference line. This finding supports our results presented in Fig 6, showing the smaller decision error probability of 0.001 for 1 vs. 8 Gy, compared to the larger decision error probability of 0.43 for 1 vs. 2 Gy. ROC curves also show that abnormalities in the p53 system can cause decision precision loss. Comparing the normal (Fig 7A) and abnormal system ROC curves (Fig 7B), we observe that the abnormal system ROC curves are closer to the 45° reference line, meaning that more erroneous decisions are made, when there is an abnormality in the system.

### Bivariate analysis: Methods for computing decision thresholds and decision error rates using *two* time point measurements in individual cells

In this section, we analyze PTEN levels of 5000 cells measured in one hour and 30 hours after the irradiation phase. Using two variables instead of one allows to study the effect of temporal dynamical changes on decision making and signaling outcomes, and paves the way for analyzing decisions based on multiple time point data. Suppose *x* and *y* represent the ln(PTEN) levels in one hour and 30 hours, respectively, after radiation. Joint Gaussian PDF for *x* and *y* can be written as [41]

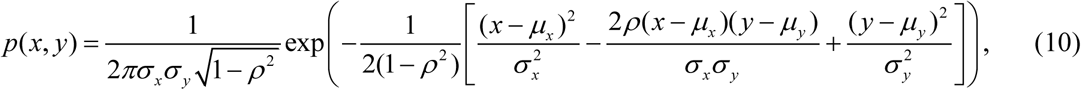

where 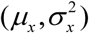 and 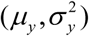 are means and variances of *x* and *y*, and *ρ* is correlation coefficient between *x* and *y*. Bivariate conditional likelihood ratio is given by *L*(*x, y*) = *p*(*x, y* | H_1_) / *p*(*x, y* | H_0_), and the optimal decision maker which minimizes the overall error probability *P*_*E*_ compares *L* (*x, y*) with the ratio *γ* = *P*(H_0_) / *P*(H_1_). The system decides H_1_ if *L*(*x, y*) > *γ*. If the hypotheses are equi-probable, i.e., *P*(H_0_) = *P*(H_1_) = 0.5, then the optimal system decides H_1_ if *p*(*x, y* | H_1_) > *p*(*x, y* | H_0_). To find the optimal decision threshold curve, we need to solve the equation *L*(*x, y*) = *γ*, i.e., *P*(H_1_) *p*(*x, y* | H_1_) = *P*(H_0_) *p*(*x, y* | H_0_), for *x* and *y*. When H_0_ and H_1_ are equi-probable, the threshold equation to be solved simplifies to *L*(*x, y*) = 1, i.e., *p*(*x, y* | H_1_) = *p*(*x, y* | H_0_). To find false alarm and miss probabilities, Equations (4) and (5) can be extended to two variables as follows:

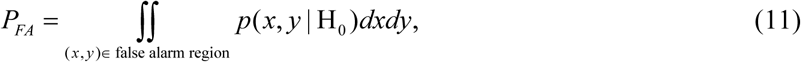

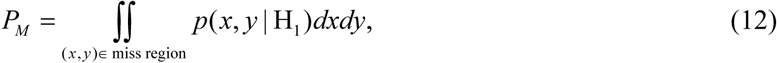

where {*x, y* : *p*(*x, y* | H_1_) > *p*(*x, y* | H_0_)} defines the false alarm region when H_0_ is true, and {*x, y* : *p*(*x, y* | H_0_) > *p*(*x, y* | H_1_)} specifies the miss region when H_1_ is true. After computing *P*_*FA*_ and *P*_*M*_, the overall decision error probability *P*_*E*_ can be calculated using Equation (3).

As an example, here we focus on Fig 8A, where two bivariate Gaussian PDFs are shown for *x* = ln(PTEN at the1^st^ hour) and *y* = ln(PTEN at the 30^th^ hour), logarithms of PTEN levels in the two data sets of IR = 1 Gy and IR = 2 Gy, with each data set consisting of 5000 cells. The mean and variance parameters of each bivariate response PDF are estimated from the associated data set. The overlap between the two bivariate PDFs in response to IR = 1 Gy and IR = 2 Gy can be better seen in the top view shown in Fig 8B. This figure also demonstrates that the decision threshold between the two PDFs is going to be a curve in the *x*-*y* plane, where the two PDFs intersect. Equation for this optimal threshold curve which minimizes the total decision error probability is given by *L*(*x, y*) = 1, where *L* is the bivariate conditional likelihood ratio defined previously. This decision threshold curve *curve*_th_ is shown together with contour plots of the two bivariate PDFs in Fig 8C. To compute the decision error probabilities using the decision threshold *curve*_th_, Equations (11) and (12) for the false alarm and miss error probabilities can be written as:

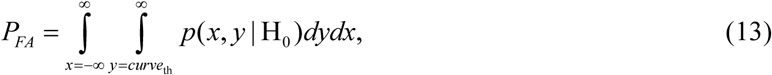

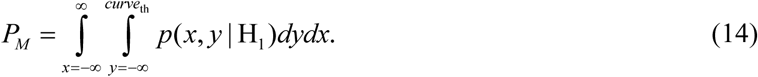

**Fig 8:**
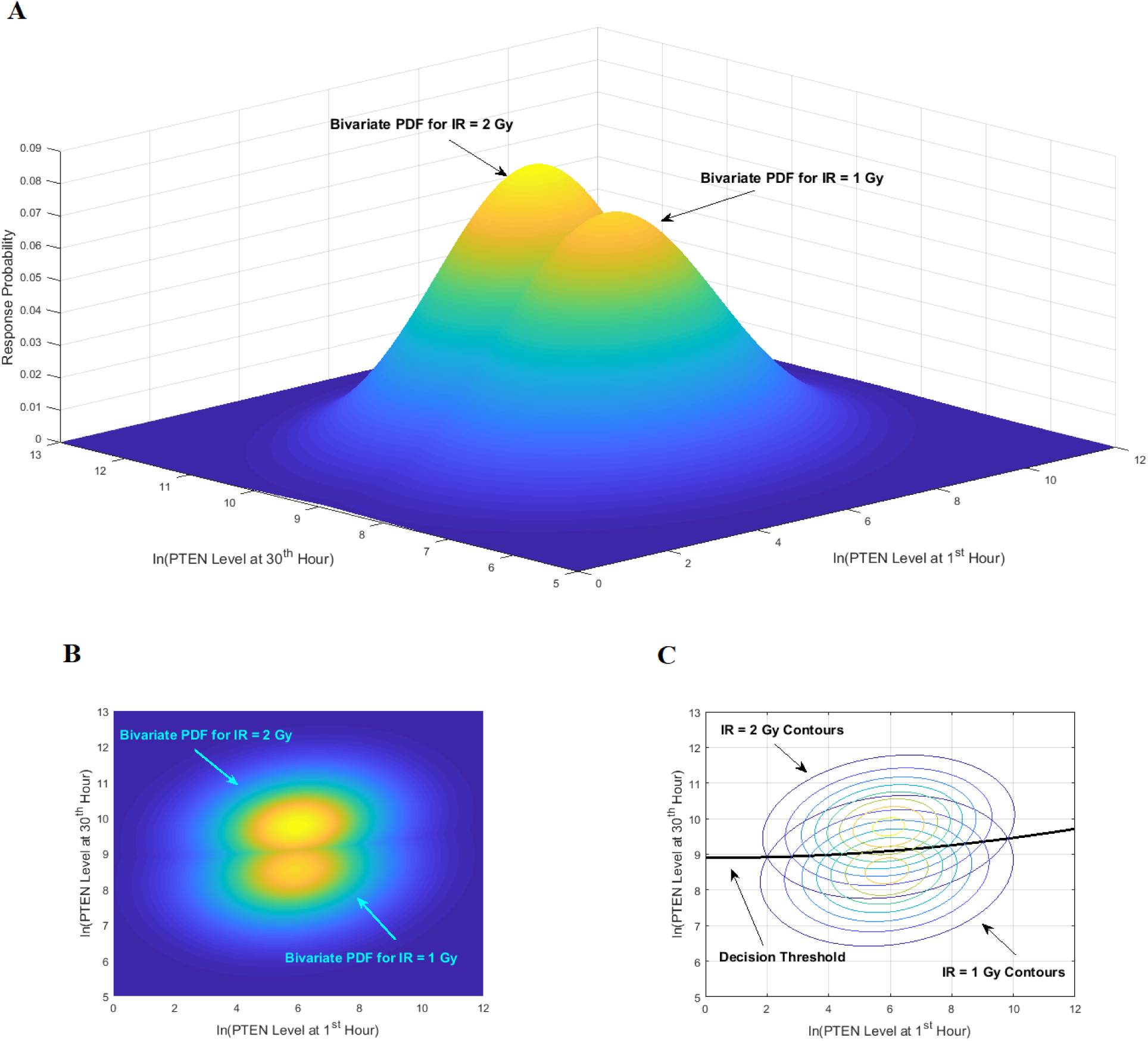
Bivariate decision making and signaling outcome analysis in the normal p53 system based on PTEN response distributions. **(A)** Bivariate Gaussian probability density functions (PDFs) for PTEN levels of cells at the 1^st^ hour and the 30^th^ hour, under IR = 1 Gy and IR = 2 Gy doses. **(B)** Top view of the two bivariate Gaussian PDFs. **(C)** Top contour view of the two bivariate Gaussian PDFs, together with the optimal maximum likelihood decision threshold curve which minimizes the total decision error probability.

After computing the integrals in Equations (13) and (14) numerically, we obtain *P*_*FA*_ = 0.24 and *P*_*M*_ = 0.26. Upon their substitution in Equation (3) and with equi-probable hypotheses, we obtain *P*_*E*_ = 0.25.

To compare the above two time point decision with individual one time point decisions, we compute decision error probabilities based on the 1^st^ hour data and the 30^th^ hour data, individually, for the IR = 1 vs. 2 Gy scenario. We obtain *P*_*E*_ = 0.5 and *P*_*E*_ = 0.27 for individual univariate decisions in one hour and 30 hours after the radiation, respectively. We observe that the bivariate decision offers significant improvement over the one hour decision, and slight improvement over the 30 hour decision. Univariate decision error probabilities at different time points are discussed in the next section, as well as how multivariate decision error probability changes, as the data of more time points are added to the decision process in a sequential manner.

### Multivariate analysis: Methods for computing decision thresholds and decision error rates using *multiple* time point measurements in individual cells

In this section, we further study the effect of system dynamics on decision making and signaling outcomes, by considering multiple time point data. More specifically, we consider PTEN levels of 5000 cells measured in 1, 10, 20, 30, 40, 50, 60 and 70 hours after the irradiation phase. Let **ω** be an *N* ×1 column vector that represents the ln(PTEN) levels at a subset or all of the aforementioned time instants. Joint Gaussian PDF for all the decision variables in **ω** can be written as [31, 40]:

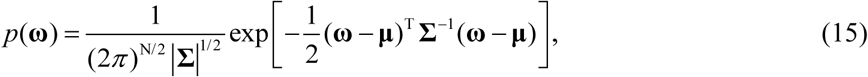

where **μ** is the *N* ×1 mean vector, **Σ** is the *N* × *N* covariance matrix, |**Σ**| and **Σ**^−1^ denote the determinant and inverse of **Σ**, respectively, and ^T^ represents matrix transpose. This multivariate Gaussian or normal PDF for the decision vector **ω** can be symbolically shown by **ω** ∼ ***𝒩*** (**μ, Σ**). For *N* = 2, Equation (15) simplifies to the bivariate PDF in Equation (10), such that:

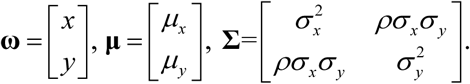

Computation of the decision error probabilities using multiple decision variables can be accomplished using discriminant functions [31, 40]:

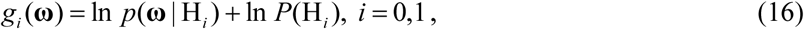

where *p*(**ω** | H_*i*_) ∼ ***𝒩*** (**μ**_*i*_, **Σ**_*i*_) and *i* is index of the discriminant function associated with the hypothesis H_*i*_. In our case we have *i* = 0,1, referring to our two hypotheses in Equation (1). For any hypothesis H_*i*_, substitution of (15) in (16) simplifies its discriminant function to:

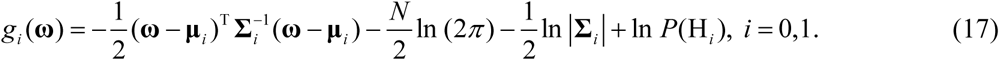

Using the discriminant functions in (17) and for a given **ω**, the optimal decision making system decides H_0_ if *g*_0_(**ω**) > *g*_1_(**ω**), and decides H_1_ if *g*_1_(**ω**) > *g*_0_(**ω**). The false alarm probability *PFA* is the probability of deciding H_1_, i.e., *g*_1_(**ω**) > *g*_0_(**ω**), whereas in fact H_0_ is true. On the other hand, the miss probability *P*_*M*_ is the probability of deciding H_0_, i.e., *g*_0_(**ω**) > *g*_1_(**ω**), although indeed H_1_ is true. Computing *P*_*FA*_ and *P*_*M*_ using multivariate PDFs directly entail multivariate integrations over regions defined by decision surfaces. Given the complexities of such computations, as a simpler alternative we calculate *P*_*FA*_ and *P*_*M*_ using the data directly, by counting the number of times that false alarm and miss event occur, respectively, after comparing the discriminant function values *g*_1_(**ω**) and *g*_0_(**ω**) for each **ω**, and then divide them by the total number of data points. The overall decision error probability *P*_*E*_ can be calculated using Equation (3). Another method for computing *P*_*FA*_ and *P*_*M*_ relies on characteristic functions [42].

#### Single-variable decision making and signaling outcome analysis as time evolves

To understand how decision making and signaling outcomes may change over time, first we look at the decision error probability *P*_*E*_ using PTEN levels measured at individual consecutive time instants (Fig 9A), for the 1 vs. 2 Gy scenario. A noteworthy observation is that the decision error exhibits a minimum value. The minimum occurs in 20 hours after the radiation. This can be visually explained by the amount of overlap of PTEN histograms at each individual time point. For instance, we provide histograms of PTEN levels at the 20^th^ and the 70^th^ hours in Fig 10, for IR = 1 and 2 Gy doses. We observe that the 20^th^ hour histograms have less overlap than the 70^th^ hour histograms, shown in Fig 10A and Fig 10B, respectively, which results in the smaller *P*_*E*_ at the 20^th^ hour in Fig 9A.

**Fig 9:**
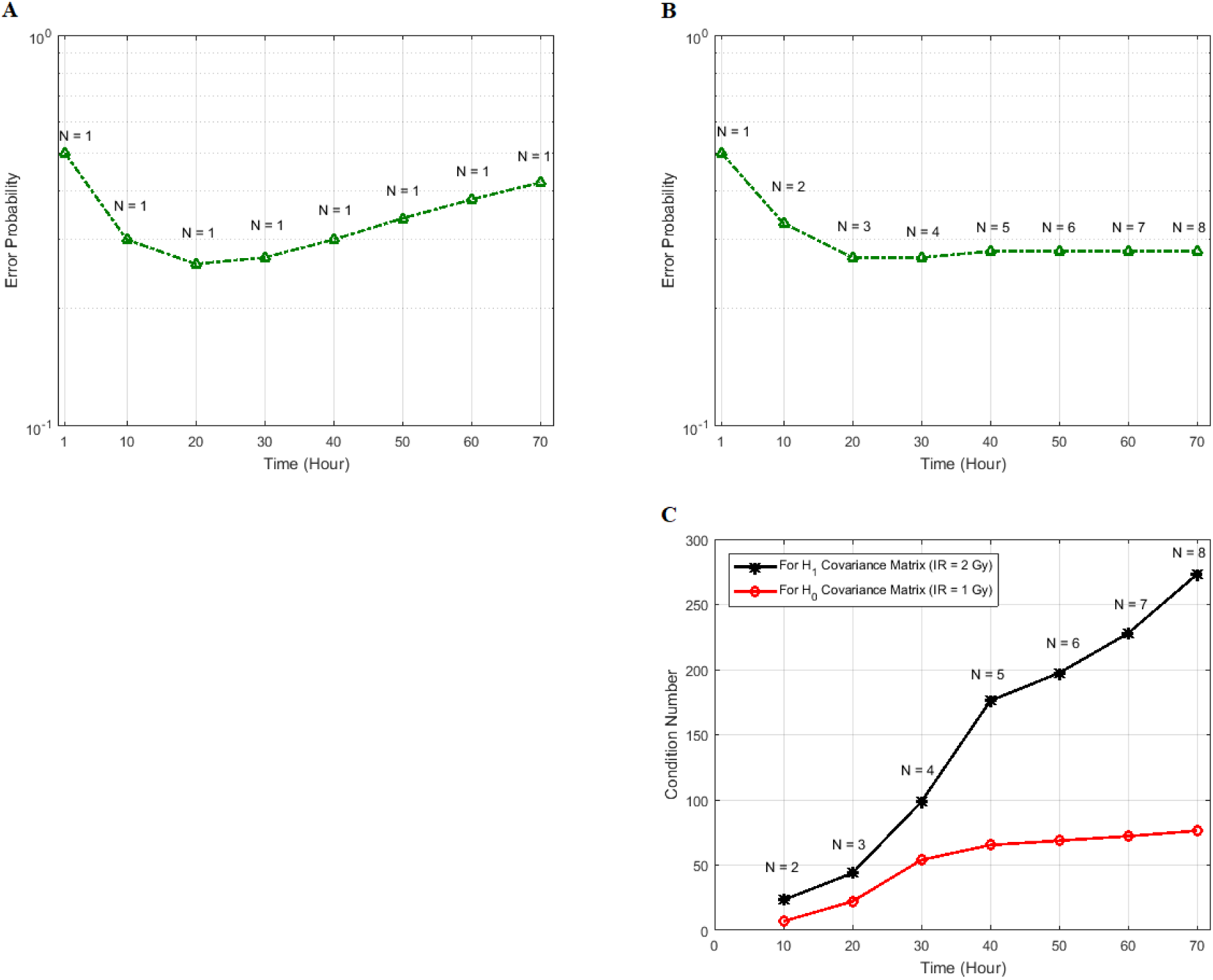
Decision error probabilities versus time in the normal p53 system: A single versus multiple time point study. **(A)** *P*_*E*_ as a function of time for the 1 vs. 2 Gy radiation scenario, computed using only the PTEN data of a single, *N* = 1, individual time instant. This assumes at any given time, decision is made based on the data of that time only. Having a minimum error probability at the 20^th^ hour is noteworthy. **(B)** *P*_*E*_ as a function of time for the 1 vs. 2 Gy radiation scenario, computed using the PTEN data of *N* time instants, *N* = 1, 2, …, 8 (*N* = 1 means the PTEN data of the 1^st^ hour, *N* = 2 refers to the PTEN data of the 1^st^ and the 10^th^ hours, *N* = 3 indicates the PTEN data of the 1^st^, the 10^th^, and the 20^th^ hours, etc.). This assumes at any given time, decision is made based on the data of that time, plus the data of the previous time instants, which means accumulating the data to make a decision. It is observed that *P*_*E*_ first decreases, and after a certain point, it remains nearly constant. **(C)** Condition numbers of **Σ** _0_ and **Σ**_1_, the *N* × *N* covariance matrices of the data for the two hypotheses H_0_ and H_1_, for IR = 1 and 2 Gy, respectively, as *N* increases from 2 to 8. When *N* increases, condition numbers of both of the covariance matrices **Σ** _0_ and **Σ**_1_ increase. On the other hand, a large condition number for a covariance matrix implies large correlations among some of its random variables. Therefore, as time evolves after a certain point, the suggested sequential decision maker incorporates a new observation that is correlated with the previously used observations. The correlation does not allow the decision error probability *P*_*E*_ to decrease beyond a certain point, although *N* constantly increases.

**Fig 10:**
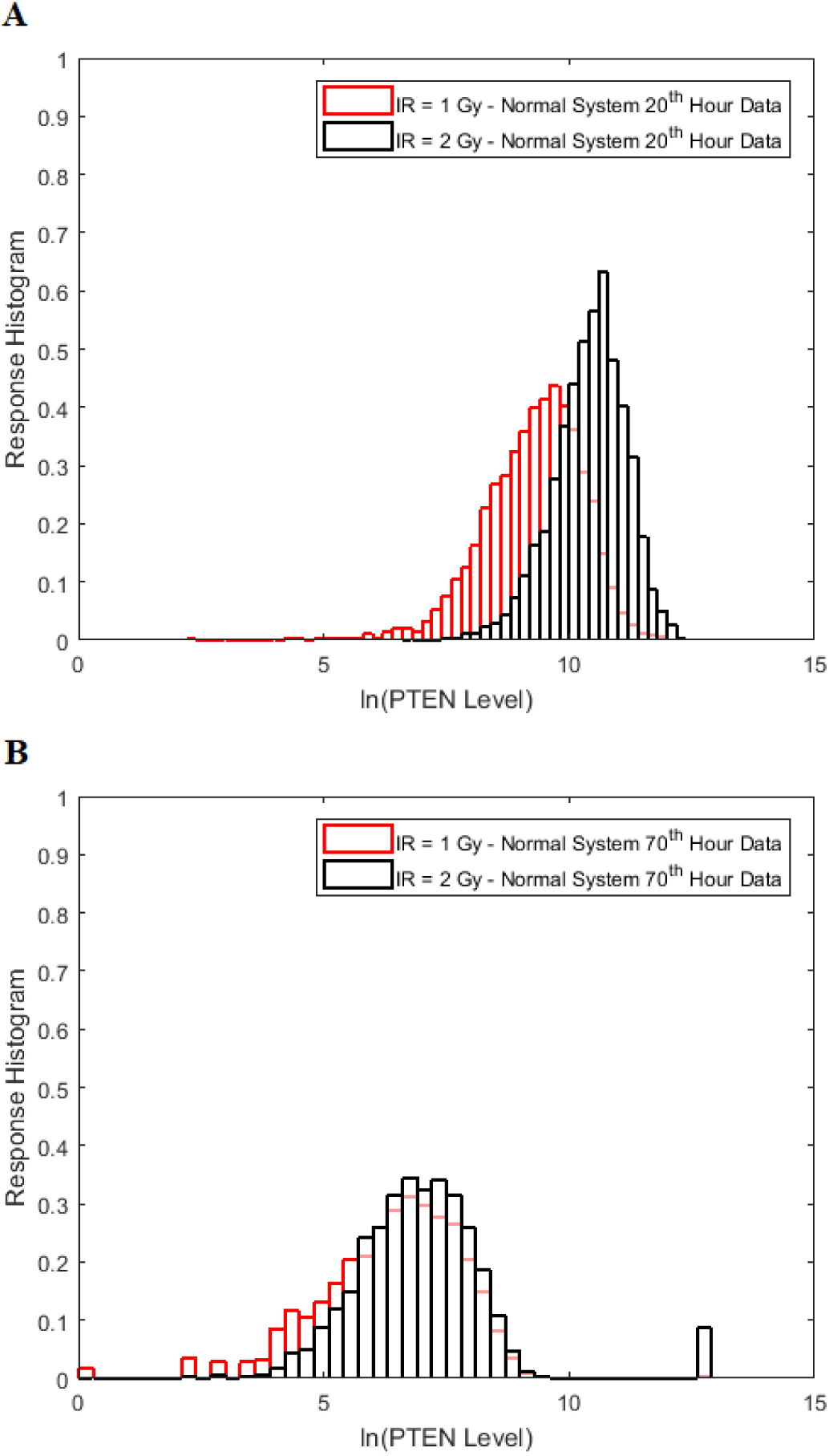
Comparison of the histograms of cells PTEN levels at the 20^th^ and the 70^th^ hours under IR = 1 Gy and 2 Gy doses in the normal p53 system. **(A)** Histograms of the 20^th^ hour PTEN data under IR = 1 and 2 Gy doses, which show less overlap. **(B)** Histograms of the 70^th^ hour PTEN data under IR = 1 and 2 Gy doses, which show more overlap.

#### Multi-variable decision making and signaling outcome analysis as time evolves

Now we focus on studying how decision making works, if data of *N* time instants are utilized, such that *N* = 1, 2, …, 8 (Fig 9B). In the figure, *N* = 1 means the PTEN data of the 1^st^ hour, *N* = 2 refers to the PTEN data of the 1^st^ and the 10^th^ hours, *N* = 3 indicates the PTEN data of the 1^st^, the 10^th^, and the 20^th^ hours, etc. This assumes at any given time, decision is made based on the data of that given time, plus the data of the previous time instants, which means progressively accumulating the data to make decisions. It is observed in Fig 9B that *P*_*E*_ first decreases, and after a certain point, it remains nearly constant. To understand this behavior, we note that if the data collected at various time instants are independent, then error probability of a decision making system that performs sequential hypothesis testing decreases, as the number of observations *N* increases [42]. This property of a multivariate sequential decision maker is intuitively appealing. However, if the data collected at various time instants are correlated, performance of the multivariate sequential decision maker can significantly degrade and its error probability does not necessarily decrease, as *N* increases [42].

To examine possible temporal correlations among the data that the suggested sequential decision strategy employs, we compute condition numbers of **Σ**_0_ and **Σ**_1_, the *N* × *N* covariance matrices of the data for the two hypotheses H_0_ and H_1_, for IR = 1 and 2 Gy, respectively, as *N* increases from 2 to 8 (Fig 9C). The condition number of a matrix is the ratio of its largest singular value to its smallest. A large condition number indicates that the matrix is nearly singular. On the other hand, a near singular covariance matrix of several random variables means that some of the random variables are highly correlated. Therefore, a large condition number for a covariance matrix implies large correlations among some of its random variables. We observe in Fig 9C that as *N* increases, condition numbers of both of the covariance matrices **Σ**_0_ and **Σ**_1_ increase. This means as time evolves after a certain point, the suggested sequential decision maker incorporates a new observation that is correlated with the previously used observations. The correlation does not allow the decision error probability to decrease beyond a certain point, although *N* constantly increases (Fig 9B).

#### Multi-variable analysis of two or more molecules: Methods for computing decision thresholds and decision error rates using their concentration measurements in individual cells

So far we have focused on multi-variable decision making and signaling outcome analysis for one molecule at different time instants. However, the introduced methods and algorithms are not limited to the outcome analyses for just one molecule, and they can be applied to various other scenarios and studies. In fact, they can be used to analyze and compute decision error rates based on concentration levels of two or more molecules, measured simultaneously or even at different time instants. For example, if decision and outcome analysis are going to be conducted based on simultaneous concentration level measurements of two molecules labeled by *x* and *y*, then Equations (10)-(14) can be used to find the maximum likelihood bivariate decision strategy and its minimum error probability. As a more elaborate example, suppose concentration levels of molecule A measured at time instants *t*_1_ and *t*_2_ are labeled as variables *x* and *y*, respectively, concentration levels of molecule B measured at *t*_1_ and *t*_2_ are labeled as variables *v* and *w*, respectively, and finally concentration levels of molecule C measured at *t*_1_ and *t*_2_ are labeled as variables *Ψ* and *ζ*, respectively. The 6 ×1 decision vector **ω** including all these six decision variables can be defined as **ω** =[*x y v w Ψ ζ*]^T^, where ^T^ stands for transpose. Now Equations (15)-(17) can be used to find the maximum likelihood six-variate decision strategy and its minimum decision error probability.

### Beyond binary decisions: Ternary decisions and signaling outcomes, and ternary error probabilities

While the focus of this paper is on binary hypothesis testing, it is possible to develop a multiple hypothesis testing model for outcome analysis, where there exist more than two possible outcomes. This entails more erroneous decisions than false alarm and miss events. Optimal decision thresholds and error probabilities for all the incorrect decisions can be similarly computed. For example, assume there are three different signaling outcomes depending on concentration level of a hypothetical molecule called MOL, whose level can fall within one of three regions, which results in the following three possible hypotheses:

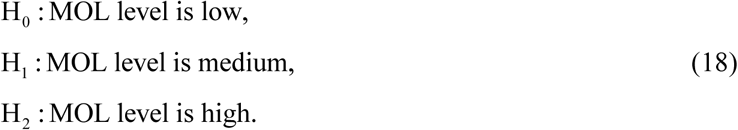

Let us assume under each condition, PDF of the MOL level represented by *x* is normal or Gaussian, i.e., *x* ∼ ***𝒩*** (*μ*_*i*_, *σ*^2^) such that *μ*_0_ < *μ*_1_ < *μ*_2_, where variances are assumed to be equal, to simplify the notation. These PDFs are shown in Fig 11, with *μ*_0_ = 5, *μ*_1_ =10, *μ*_2_ =15, and *σ*^2^ = 2.25. By extending the binary decision errors presented earlier in Equations (8) and (9), ternary decision errors for the three hypotheses can be written as:

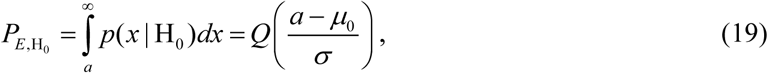

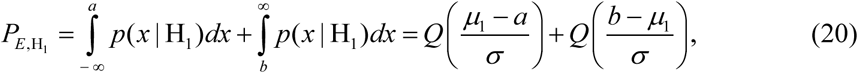

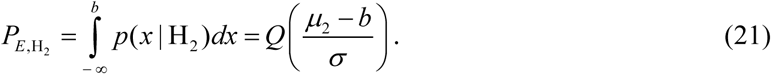

**Fig 11:**
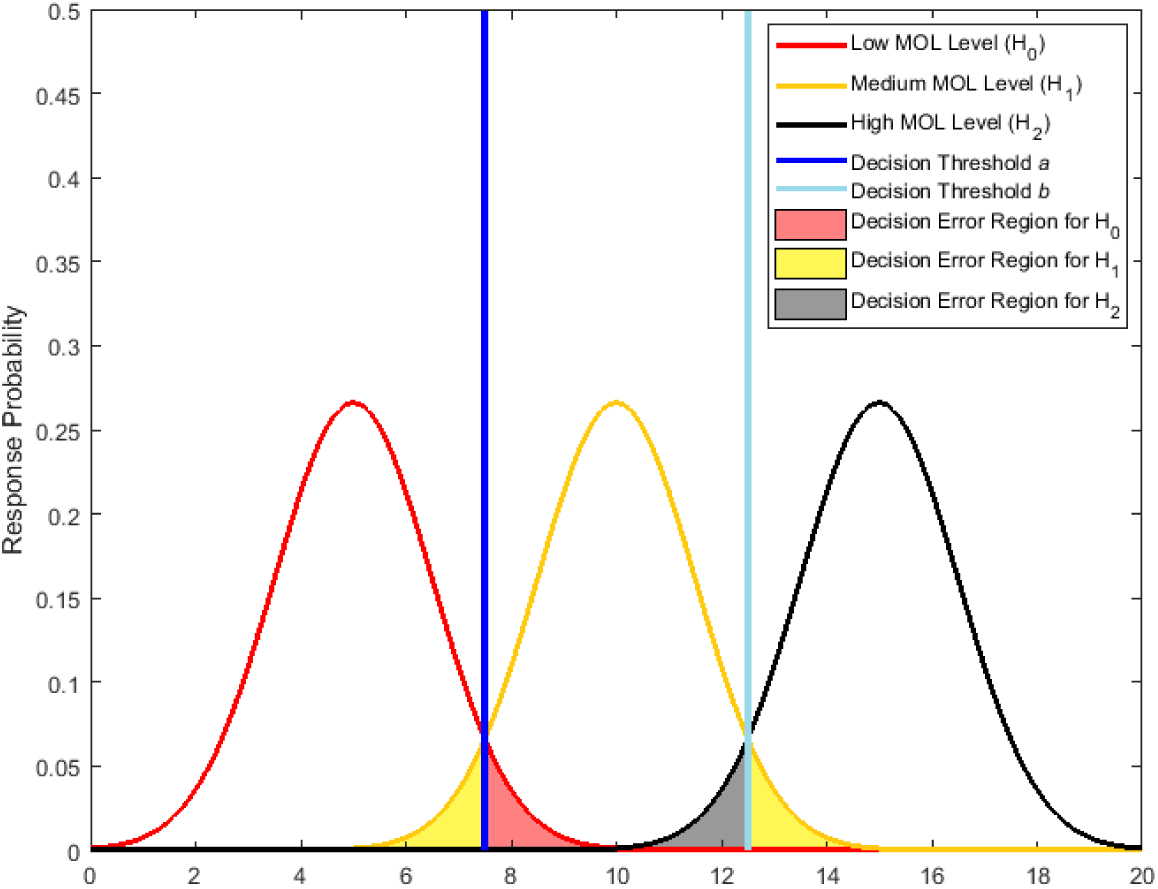
Response probability density functions of a hypothetical molecule called MOL whose level entails a ternary decision making process with three signaling outcomes. Shaded tail areas with the same color represent decision error regions associated with each specific hypothesis. Assuming equi-probable hypotheses, optimal maximum likelihood decision thresholds which minimize the total decision error probability are shown by vertical blue lines at the points of intersection of the probability density functions.

In the above equations, *a* and *b* are thresholds to decide between H_0_ and H_1_, and between H_1_ and H_2_, respectively. This means the decision regions for the three hypotheses can be written as:

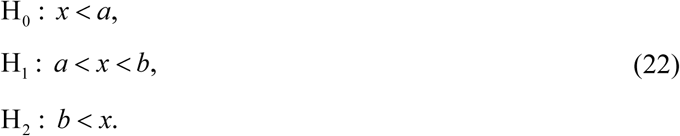

For equi-probable hypotheses and similarly to the derivation that lead to Equation (7), optimal decision thresholds which minimize the total decision error probability can be shown to be:

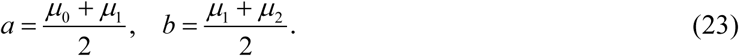

Upon substituting (23) in (19)-(21), the total error probability in making ternary decisions can be written as:

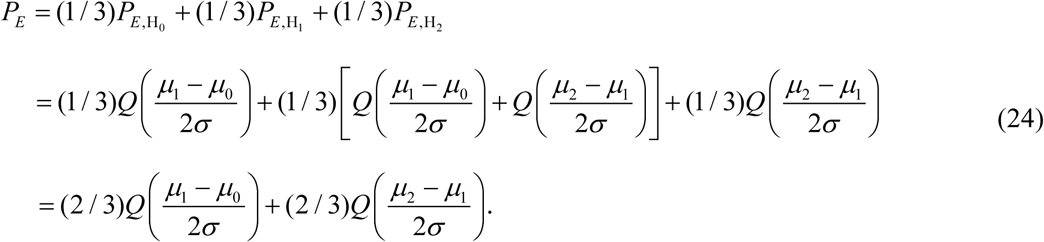

As a reference, for the binary decision making problem and outcome analysis studied earlier in the paper and using Equations (8) and (9), the total error probability in making binary decisions with equal variances simplifies to:

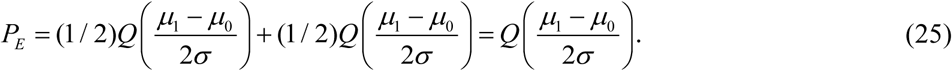

To compare ternary and binary error probabilities, let us assume *μ*_2_ − *μ*_1_ = *μ*_1_ − *μ*_0_ = *γ*, which reduces Equations (24) and (25) to (4 / 3)*Q*(*γ* / (2*σ*)) and *Q*(*γ* / (2*σ*)), respectively. This indicates that the ternary decision error rate can be higher than the binary decision error rate, under the assumed conditions.

### On the costs of correct and incorrect decisions

In decision theory, there can be some costs associated with correct or incorrect decisions. Let *C*_*ij*_ be the cost of deciding H_*i*_ when H _*j*_ is true. To minimize the expected cost, *C*_00_ *P*(H_0_) + *C*_01_*P*(H_1_) *P*_*M*_ +*C*_10_ *P*(H_0_) *P*_*FA*_ + *C*_11_*P*(H_1_), the decision making system decides H_1_ if [30]:

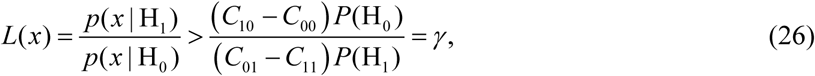

where *C*_10_ > *C*_00_ and *C*_01_ > *C*_11_. Usually the costs associated with correct decisions are zero, i.e., *C*_00_ = *C*_11_ = 0. Additionally, if there is no preference in assigning different costs to different incorrect decisions, one can choose *C*_10_ = *C*_01_. This is what we would consider as well, since we do not have a knowledge of the costs of incorrect decisions in the studied cellular system. Upon substituting *C*_00_ = *C*_11_ = 0 and *C*_10_ = *C*_01_ in the above equation, it simplifies to the following equation, which is the optimal maximum likelihood decision rule presented earlier in the paper:

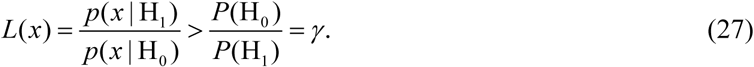

## Conclusion

This study presents a set of decision-theoretic and statistical signal processing methods and metrics for modeling and measurement of decision making processes and signaling outcomes under normal and abnormal conditions, and in the presence of noise and other uncertainties. Due to the noise, signaling malfunctions, or other factors, cells may respond differently to the same input signal. Some of these responses can be erroneous and unexpected. Here we present univariate and multivariate models and methods for decision making processes and signaling outcome analyses and as an example, apply them to an important system that is involved in cell survival and death, i.e., the p53 system shown in Fig 1 (another decision analysis example can be found in the paper by Habibi *et al.* [1]). The p53 system becomes active due to DNA damage caused by ionizing radiation (IR), and as a result, cell can take two different actions: it can either survive by repairing the DNA or trigger apoptosis. In this context, we model decisions and signaling outcomes triggered by the p53 system as a binary hypothesis testing problem, where two hypotheses are introduced in Equation (1). Regarding these two hypotheses, our approach identifies that there can be two types of incorrect decisions: *false alarm* and *miss*. To compute the likelihood of these decisions, we employ the simulator of Hat *et al.* [9], to obtain single cell data of the p53 system, by exposing the cells to different radiation doses. We consider PTEN levels in cells as the decision variable, since it is a good predictor of cell fate [9]. Our analysis focuses on low radiation dose versus high radiation dose scenarios, where we fix the low IR dose at 1 Gy, whereas we set the high IR dose at 2 Gy, 3 Gy, 4 Gy, 5 Gy, 6 Gy, 7 Gy and 8 Gy. We also analyze decision making events and signaling outcomes when an abnormality is present in the p53 system.

The incorrect decision probabilities provided in Equation (2) and the overall decision error probability in Equation (3) are computed after determining an optimal decision threshold. We obtain this decision threshold using the maximum likelihood principle which states that the best decision can be made by selecting the hypothesis that has the maximum probability of occurrence. We compute decision threshold and error probabilities using single time point data of PTEN levels in both normal and abnormal p53 systems. For 1 Gy vs. 2 Gy and 1 Gy vs. 8 Gy case studies, we present histograms, response distributions, decision thresholds, and false alarm and miss decision regions in normal and abnormal p53 systems in Fig 3 and Fig 5, respectively. Our decision analysis reveals and quantifies that more erroneous decisions are made when deciding between two nearly the same radiation doses in the normal p53 system (Fig 6). On the other hand, the difference between responses is easily identifiable for very low versus very high IR doses. This feature seems not be present in the abnormal p53 systems (Fig 6), according to our decision modeling approach. Our decision and outcome analyses and observations are further visualized and confirmed by using the receiver operating characteristic (ROC) curves (Fig 7), which are useful graphical tools to study the performance of decision making systems. We would like to note that these observations are specifically made based on the low versus high IR case studies, e.g. d_0_ vs d_1_ IRs introduced in the paper for the p53 system, as an example of a signaling network, in which the low IR dose is fixed to 1 Gy (d_0_ = 1 Gy) and the high IR dose is ranging from 2 Gy up to 8 Gy (d_1_ = 2, 3, …, 8 Gy). Such conclusions may not be generalized to other biological hypotheses and systems, while the proposed framework and its analytical tools, whose introduction has been the main goal of this paper, can be similarly used.

In addition to the above univariate single time point analysis, we extend our signaling outcome modeling framework to dynamical multi-time point measurements and multi-dimensional decision making algorithms, to see how the number of decision variables affects the decisions and signaling outcomes over time. To introduce the concepts, first we conduct a bivariate analysis, for which bivariate response distributions of cells PTEN levels measured at two different time instants are shown in Fig 8, as well as the optimal maximum likelihood decision boundary. Then we introduce a multivariate dynamic decision modeling framework, for the general scenario where there are more than two decision variables over time. This allows to model and understand how decision error probability changes over time, if at any time the decision is made based on the current observation, together with the previous observations. We observe in Fig 9B that as the decision making strategy incorporates more and more PTEN data of various time instants into its decisions, for the p53 system exposed to two radiation doses of 1 and 2 Gy, the decision error probability reaches its smallest value at a certain time instant. However, adding more data afterwards does not necessarily improve the decision precision, i.e., the decision error probability does not necessarily decrease as *N* increases with time (Fig 9B). We show that this behavior can be related to the correlations that exist among the PTEN levels measured at different times (Fig 9C).

Although we focus on multi-variable decision making and signaling outcome analysis for one molecule at different time instants, the introduced methods and algorithms are not limited to the outcome analyses for just one molecule. They can be applied to various other scenarios and studies. For instance, they can be used to analyze decision strategies and compute decision error rates based on concentration levels of two or more molecules, measured simultaneously or even at different time instants.

We finally show how the introduced binary decision making and signaling outcome analysis models can be extended to more than two decisions, i.e., more than two hypotheses. A ternary scenario with three signaling outcomes is analyzed as an example, and it is shown that under certain conditions, ternary decision error probability can be higher than the binary one.

The methods and models presented here can be expanded to describe the performance and precision of more complex systems and networks such as the ones whose inputs are multiple ligands or secondary messengers and whose outputs are several transcription factors involved in certain cellular functions. Analyzing concentration levels of these transcription factors over time using the proposed approaches can model various decisions and signaling outcomes, and their probabilities, in the presence of noise or some cellular abnormalities, and in response to the input signals.

Overall, these decision-theoretic models and signaling outcome analysis methods can be beneficial for better understanding of transition from physiological to pathological conditions such as inflammatory diseases, various cancers and autoimmune diseases.

## Acknowledgements

The authors declare that no conflict of interests exist. TL was supported by the National Science Center (Poland), grant 2014/14/M/NZ6/00537. AL was supported by NIH grants: GM072024 and GM123011.

## Author Contributions

MO developed the methods, performed the simulations and computations, and wrote the manuscript.

TL contributed to and discussed the research topics, and edited the manuscript.

AL contributed to and discussed the research topics, and edited the manuscript.

ESE contributed to and discussed the research topics, and edited the manuscript.

AA conceived the ideas and concepts, developed the methods, and wrote the manuscript.

## References

1. Habibi I, Cheong R, Lipniacki T, Levchenko A, Emamian ES, Abdi A (2017) Computation and measurement of cell decision making errors using single cell data. PLoS Comput Biol 13(4): e1005436. https://doi.org/10.1371/journal.pcbi.1005436

2. Kolitz SE, Lauffenburger DA (2012) Measurement and modeling of signaling at the single-cell level. Biochemistry 51(38): 7433–7443. https://doi.org/10.1021/bi300846p PMID: 22954137

3. Cheong R, Rhee A, Wang CJ, Nemenman I, Levchenko A (2011) Information transduction capacity of noisy biochemical signaling networks. Science; 334(6054): 354–358. https://doi.org/10.1126/science.1204553 PMID: 21921160

4. Balazsi G, van Oudenaarden A, Collins JJ (2011) Cellular decision making and biological noise: From microbes to mammals. Cell; 144(6): 910–925.

5. Vousden KH, Prives C (2009) Blinded by the Light: The Growing Complexity of p53. Cell.; 137: 413-431. doi:10.1016/j.cell.2009.04.037 PMID: 19410540

6. Levine AJ (1997) p53, the cellular gatekeeper for growth and division. Cell; 88: 323-331. PMID: 9039259

7. Kastan MB, Onyekwere O, Sidransky D, Vogelstein B, Craig RW (1991) Participation of p53 protein in the cellular response to DNA damage. Cancer Res.; 51: 6304-6311. PMID: 1933891

8. Elmore S (2007) Apoptosis: A Review of Programmed Cell Death. Toxicol Pathol.; 35: 495–516. doi: 10.1080/01926230701320337 PMID: 17562483

9. Hat B, Kochanczyk M, Bogdal MN, Lipniacki T (2016) Feedbacks, Bifurcations, and Cell Fate Decision-Making in the p53 System. PLoS Comput Biol 12(2): e1004787. https://doi.org/10.1371/journal.pcbi.1004787

10. Siliciano JD, Canman CE, Taya Y, Sakaguchi K, Appella E, Kastan MB (1997) DNA damage induces phosphorylation of the amino terminus of p53. Genes Dev.; 11: 3471-3481. PMID: 9407038

11. Vogelstein B, Lane D, Levine AJ (2000) Surfing the p53 network. Nature; 408: 307–310.

12. Rothkamm K, Krüger I, Thompson LH, Löbrich M (2003) Pathways of DNA Double-Strand Break Repair during the Mammalian Cell Cycle. Mol Cell Biol; 23: 5706–5715.

13. Alberts B, Johnson A, Lewis J, Raff M, Roberts K, Walter P (2002) Molecular Biology of the Cell. 4th ed. Garland Science.

14. Iliakis G (1991) The role of DNA double strand breaks in ionizing radiation-induced killing of eukaryotic cells. BioEssays; 13: 641–648. doi: 10.1002/bies.950131204 PMID: 1789781

15. Vilenchik MM, Knudson AG (2003) Endogenous DNA double-strand breaks: Production, fidelity of repair, and induction of cancer. Proc Natl Acad Sci USA; 100: 12871–12876. doi: 10.1073/pnas.2135498100 PMID: 14566050

16. Saito S, Goodarzi AA, Higashimoto Y, Noda Y, Lees-Miller SP, Appella E, et al. (2002) ATM Mediates Phosphorylation at Multiple p53 Sites, Including Ser46, in Response to Ionizing Radiation. J Biol Chem.; 277: 12491-12494. PMID: 11875057

17. Bakkenist CJ, Kastan MB (2003) DNA damage activates ATM through intermolecular autophosphorylation and dimer dissociation. Nature.; 421: 499–506. doi: 10.1038/nature01368 PMID: 12556884

18. Maya R, Balass M, Kim S-T, Shkedy D, Leal J-FM, Shifman O, et al. (2001) ATM-dependent phosphorylation of Mdm2 on serine 395: role in p53 activation by DNA damage. Genes Dev.; 15: 1067–1077. doi: 10.1101/gad.886901 PMID: 11331603

19. Shieh SY, Taya Y, Prives C (1999) DNA damage-inducible phosphorylation of p53 at N-terminal sites including a novel site, Ser20, requires tetramerization. EMBO J.; 18: 1815–1823. doi: 10.1093/emboj/18.7.1815 PMID: 10202145

20. Banin S, Moyal L, Shieh S, Taya Y, Anderson CW, Chessa L, et al. (1998) Enhanced phosphorylation of p53 by ATM in response to DNA damage. Science; 281: 1674-1677. PMID: 9733514

21. Canman CE, Lim D-S, Cimprich KA, Taya Y, Tamai K, Sakaguchi K, et al. (1998) Activation of the ATM Kinase by Ionizing Radiation and Phosphorylation of p53. Science; 281: 1677–1679. doi: 10.1126/science.281.5383.1677 PMID: 9733515

22. Barak Y, Juven T, Haffner R, Oren M (1993) mdm2 expression is induced by wild type p53 activity. EMBO J.; 12: 461-468. PMID: 8440237

23. Fiscella M, Zhang H, Fan S, Sakaguchi K, Shen S, Mercer WE, et al. (1997) Wip1, a novel human protein phosphatase that is induced in response to ionizing radiation in a p53-dependent manner. Proc Natl Acad Sci USA; 94: 6048-6053. PMID: 9177166

24. Choi J, Nannenga B, Demidov ON, Bulavin DV, Cooney A, Brayton C, et al. (2002) Mice deficient for the wildtype p53-induced phosphatase gene (Wip1) exhibit defects in reproductive organs, immune function, and cell cycle control. Mol Cell Biol.; 22: 1094-1105. PMID: 11809801

25. Shreeram S, Hee WK, Demidov ON, Kek C, Yamaguchi H, Fornace AJ, et al. (2006) Regulation of ATM/p53-dependent suppression of myc-induced lymphomas by Wip1 phosphatase. J Exp Med.; 203: 2793–2799. doi: 10.1084/jem.20061563 PMID: 17158963

26. Takekawa M, Adachi M, Nakahata A, Nakayama I, Itoh F, Tsukuda H, et al. (2000) p53-inducible Wip1 phosphatase mediates a negative feedback regulation of p38 MAPK-p53 signaling in response to UV radiation. EMBO J.; 19: 6517–6526. doi: 10.1093/emboj/19.23.6517 PMID: 11101524

27. Stambolic V, MacPherson D, Sas D, Lin Y, Snow B, Jang Y, et al. (2001) Regulation of PTEN transcription by p53. Mol Cell.; 8: 317-325. PMID: 11545734

28. Hlobilkova A, Knillova J, Svachova M, Skypalova P, Krystof V, Kolar Z (2006) Tumour suppressor PTEN regulates cell cycle and protein kinase B/Akt pathway in breast cancer cells. Anticancer Res.; 26: 1015-1022. PMID: 16619501

29. Bogdal MN, Hat B, Kochanczyk M, Lipniacki T (2013) Levels of pro-apoptotic regulator Bad and anti-apoptotic regulator Bcl-xL determine the type of the apoptotic logic gate. BMC Syst Biol.; 7: 67. doi: 10.1186/1752-0509-7-67 PMID: 23883471

30. Kay SM (1998) Fundamentals of Statistical Signal Processing: Detection Theory. PTR Prentice-Hall.

31. Duda RO, Hart PE and Stork DG (2001) Pattern Classification. John Wiley & Sons.

32. Lu X, Ma O, Nguyen T-A, Jones SN, Oren M, Donehower LA (2007) The Wip1 Phosphatase Acts as a Gatekeeper in the p53-Mdm2 Autoregulatory Loop. Cancer Cell; 12: 342–354. doi: 10.1016/j.ccr.2007.08.033 PMID: 17936559

33. Li J, Yang Y, Peng Y, Austin RJ, van Eyndhoven WG, Nguyen KCQ, et al. (2002) Oncogenic properties of PPM1D located within a breast cancer amplification epicenter at 17q23. Nat Genet.; 31: 133–134. doi: 10.1038/ng888 PMID: 12021784

34. Saito-Ohara F, Imoto I, Inoue J, Hosoi H, Nakagawara A, Sugimoto T, et al. (2003) PPM1D is a potential target for 17q gain in neuroblastoma. Cancer Res.; 63: 1876-1883. PMID: 12702577

35. Bulavin DV, Demidov ON, Saito S, Kauraniemi P, Phillips C, Amundson SA, et al. (2002) Amplification of PPM1D in human tumors abrogates p53 tumor-suppressor activity. Nat Genet.; 31: 210–215. doi: 10.1038/ng894 PMID: 12021785

36. Castellino RC, Bortoli MD, Lu X, Moon S-H, Nguyen T-A, Shepard MA, et al. (2007) Medulloblastomas overexpress the p53-inactivating oncogene WIP1/PPM1D. J Neurooncol.; 86: 245–256. doi: 10.1007/s11060-007-9470-8 PMID: 17932621

37. Hirasawa A, Saito-Ohara F, Inoue J, Aoki D, Susumu N, Yokoyama T, et al. (2003) Association of 17q21-q24 Gain in Ovarian Clear Cell Adenocarcinomas with Poor Prognosis and Identification of PPM1D and APPBP2 as Likely Amplification Targets. Clin Cancer Res.; 9: 1995-2004. PMID: 12796361

38. Rauta J, Alarmo E-L, Kauraniemi P, Karhu R, Kuukasjärvi T, Kallioniemi A (2006) The serine-threonine protein phosphatase PPM1D is frequently activated through amplification in aggressive primary breast tumours. Breast Cancer Res Treat.; 95: 257–263. doi: 10.1007/s10549-005-9017-7 PMID: 16254685

39. Geva-Zatorsky N, Rosenfeld N, Itzkovitz S, Milo R, Sigal A, Dekel E, et al. (2006) Oscillations and variability in the p53 system. Mol Syst Biol.; 2:2006.0033

40. Van Trees HL, Bell KL, Tian Z (2013) Detection, Estimation and Modulation Theory, Part I: Detection, Estimation, and Filtering Theory. 2nd ed. Wiley.

41. Papoulis A (1991) Probability, Random Variables, and Stochastic Processes. 3rd ed. McGraw-Hill.

42. Fukunaga K (1990) Introduction to Statistical Pattern Recognition. 2nd ed. Academic Press.

